# Single cell analysis of Barrett’s esophagus and carcinoma reveals cell types conferring risk via genetic predisposition

**DOI:** 10.1101/2024.11.18.624112

**Authors:** Marten C. Wenzel, Ann-Sophie Giel, Patrick S. Plum, Sascha Hoppe, Marek Franitza, Christoph Jonas, Pouria Dasmeh, René Thieme, Yue Zhao, Dominik Heider, Claire Palles, Rebecca Claire Fitzgerald, Christiane J. Bruns, Reinhard Buettner, Alexander Quaas, Ines Gockel, Carlo Maj, Seung-Hun Chon, Johannes Schumacher, Axel M. Hillmer

## Abstract

Inherited genetic variants contribute to Barrett’s esophagus (BE) and esophageal adenocarcinoma (EAC) but it is unknown which cell types are involved in this process. We performed single cell RNA-sequencing of BE, EAC and paired normal tissues and integrated data of a genome-wide association study to determine cell type-specific genetic risk and cellular processes that contribute to BE and EAC. The analysis revealed that EAC development is driven to a greater extent by local cellular processes than BE development. One cell type of BE origin (BE-EAC) and cellular processes that control the differentiation of columnar cells are of particular relevance for EAC development. Further, specific subtypes of fibroblasts and endothelial cells contribute to BE and EAC development, while dendritic cells and CD4+ memory T cells contribute exclusively to BE development. The diagnostic use of markers characterizing the identified cell types and cellular processes should be explored in future for EAC prediction.

## Background

Esophageal adenocarcinoma (EAC) is often preceded by Barrett’s esophagus (BE), a benign metaplasia of the epithelium increasing the risk of developing EAC 10- to 50-fold ^1,2^. BE is associated with gastro-esophageal reflux disease (GERD) and is likely connected to a chronically inflamed environment ^3^.

BE can undergo histological changes ranging from low- to high-grade dysplasia. At an approximate rate of 6 per 100 patient-years, patients progress during the first years of follow-up from high-grade dysplasia to EAC, which has a devastating 5-year survival rate ^4,5^. The incidence of EAC increased dramatically over the last decades, ^6^ emphasizing the need to better understand the cellular processes that determine which BE patients progress to EAC.

Most single cell RNA-sequencing (scRNA-seq) studies in the context of BE and EAC focused on the in-detail description of BE in order to investigate the cell of origin of BE. While the cell of origin of BE has been investigated by Owen et al. ^7^ and Nowicki-Osuch et al. ^8^, our understanding of which other cell types beyond the metaplastic cells contribute to EAC progression is limited.

In the present study, we performed scRNA-seq of normal esophagus (EN) and gastric fundus (GFN) of nine patients. Five of them were BE patients of whom in addition to the two normal tissue types also BE tissues were analyzed and four patients had EAC of whom EAC tissues were analyzed in addition to the normal tissues. We integrated our scRNA-seq data and data of a genome-wide association study (GWAS) of BE/EAC ^9^ to determine which cell types show enrichment for the expression of genes that are located at BE and/or EAC risk loci. We identify specific cell types and cellular processes that contribute to BE and to EAC progression based on germline genetic risk.

## Results

### Single cell RNA-sequencing of BE, EAC and normal tissues

To characterize cell types of BE and EAC and compare the cell type features with their counterparts of the same patient’s EN and GFN we collected biopsies of BE patients and tissue samples of EAC patients (**Supplementary Table 1**). Tissue samples were collected in a standardized manner regarding the distances of sample sites, then dissociated into single cells and sequenced for their RNA content using gel bead-in-emulsion (GEM) technology (10x Genomics, **Figure 1A**).

**Figure 1.**
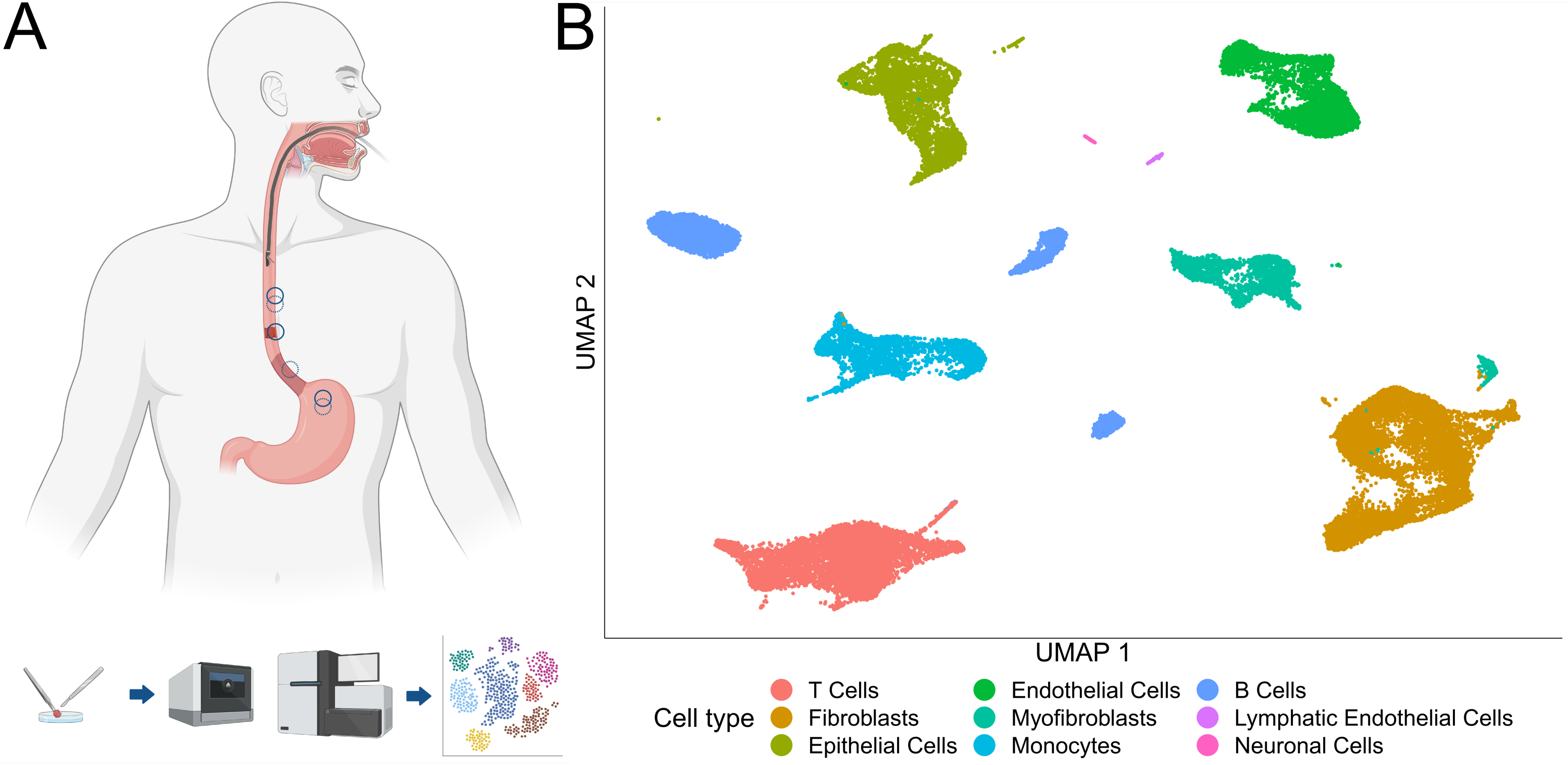
Single cell RNA-sequencing of biopsies of the esophagus and gastric fundus. **A** Graphical description of the experimental setup. Samples were taken from healthy esophageal and stomach tissue of every EAC and BE patient, respectively. Depending on the diagnosis, either samples of EAC or BE were taken as a third sample of each patient. All three tissue types per patient were dissociated separately. Single cells were prepared for fluorescence activated cell sorting (FACS) and RNA-sequencing using the 10x Genomics Chromium technology with one library per sample (three per patient). The data were analyzed using the Seurat platform, among others. Light red esophagus region close to the stomach represents BE, dark red region represents EAC. Solid line circles indicate sampling strategy for EAC and dashed line circles for BE patients. **B** UMAP representation of the whole data set. Each data point represents a single cell with coloring according to initial cell type family annotation.

After quality control and clustering, we used the expression of typical cell type markers to assign preliminary cell type labels (**Figure 1B, Supplementary Figures 1-3**). The group of epithelial cells could be identified by the expression of the marker gene *EPCAM*. The associated clusters contained cells from BE and EAC, but also from EN and GFN (**Supplementary Figures 2, 3**). We focus in the following on the epithelial cells. For characterization of mesenchymal and immune cells, see **Supplementary Note 1 and Supplementary Figure 4**.

To get a better understanding of the features of the epithelial cells we used the clusters identified as epithelial cells to run a separate analysis (**Figure 2A-D**). After initial unsupervised clustering, differences between the analyzed clusters were examined which justified merging single unsupervised clusters to functionally more coherent groups (**Supplementary Figure 5A-F, Supplementary Table 2**) allowing the identification of generic cell types based on top differentially expressed genes.

**Figure 2.**
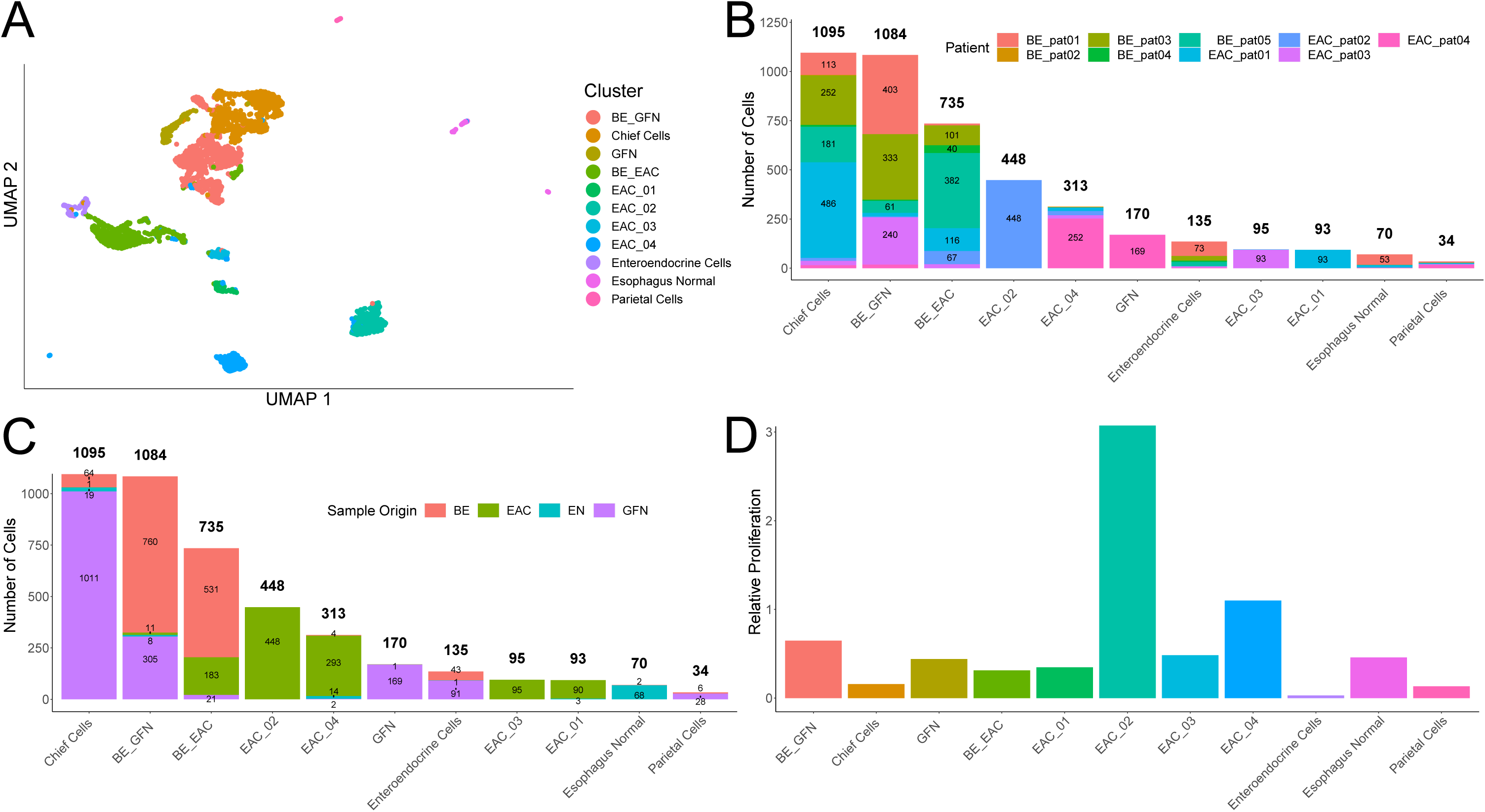
Epithelial cells show similarities between BE and GFN cells and dissimilarities between EAC cells of different patients. **A** UMAP representation of the epithelial clusters. Notably, the tumor clusters lie apart from each other with a high degree of homogeneity regarding patient origin. **B** Composition of epithelial clusters regarding patient origin. **C** Composition of epithelial clusters regarding sample origin. **D** Clusters EAC-02 and EAC-04 show the highest relative proliferativeness.

The clusters later annotated as cancer cells, were found to be largely homogenous regarding patient origin in contrast to the other clusters (**Figure 2B**). In addition, epithelial cells of BE and GFN origin showed similarities across patients (BE-GFN and enteroendocrine cells) as well as epithelial cells of BE and EAC origin (BE-EAC, **Figure 2C**).

Following the strategy presented by Tirosh et al. ^10^, we determined a cell cycle phase, i.e. G1, S, and G2/M (**Supplementary Figure 5G**), and calculated a relative proliferation index by dividing the number of cells in S and G2/M phase by the number of cells in G1 within a given cluster. EAC-02 and EAC-04 had the highest relative proliferativeness among all epithelial clusters followed by BE-GFN (**Figure 2D**).

### Characterization of epithelial cells and detection of typical genomic aberrations in EAC

Next, we annotated the epithelial clusters as chief cells, parietal cells, esophagus normal and enteroendocrine cells based on the expression of marker genes (**Figure 3A** and **Supplementary Note 2, Supplementary Tables 3 and 4**). *TFF1*, *TFF2*, and *MUC5AC* were highly expressed in clusters BE-GFN, GFN, and BE-EAC as well as in cluster EAC-01. In addition, *MSMB* was highly expressed in clusters BE-GFN and GFN, but almost absent in cancer cell containing clusters, including BE-EAC. EAC-02 and weaker EAC-04 almost exclusively expressed *WNT11*. *CEACAM6* was present in all cancer clusters, including BE-EAC, with highest expression in EAC-01.

**Figure 3.**
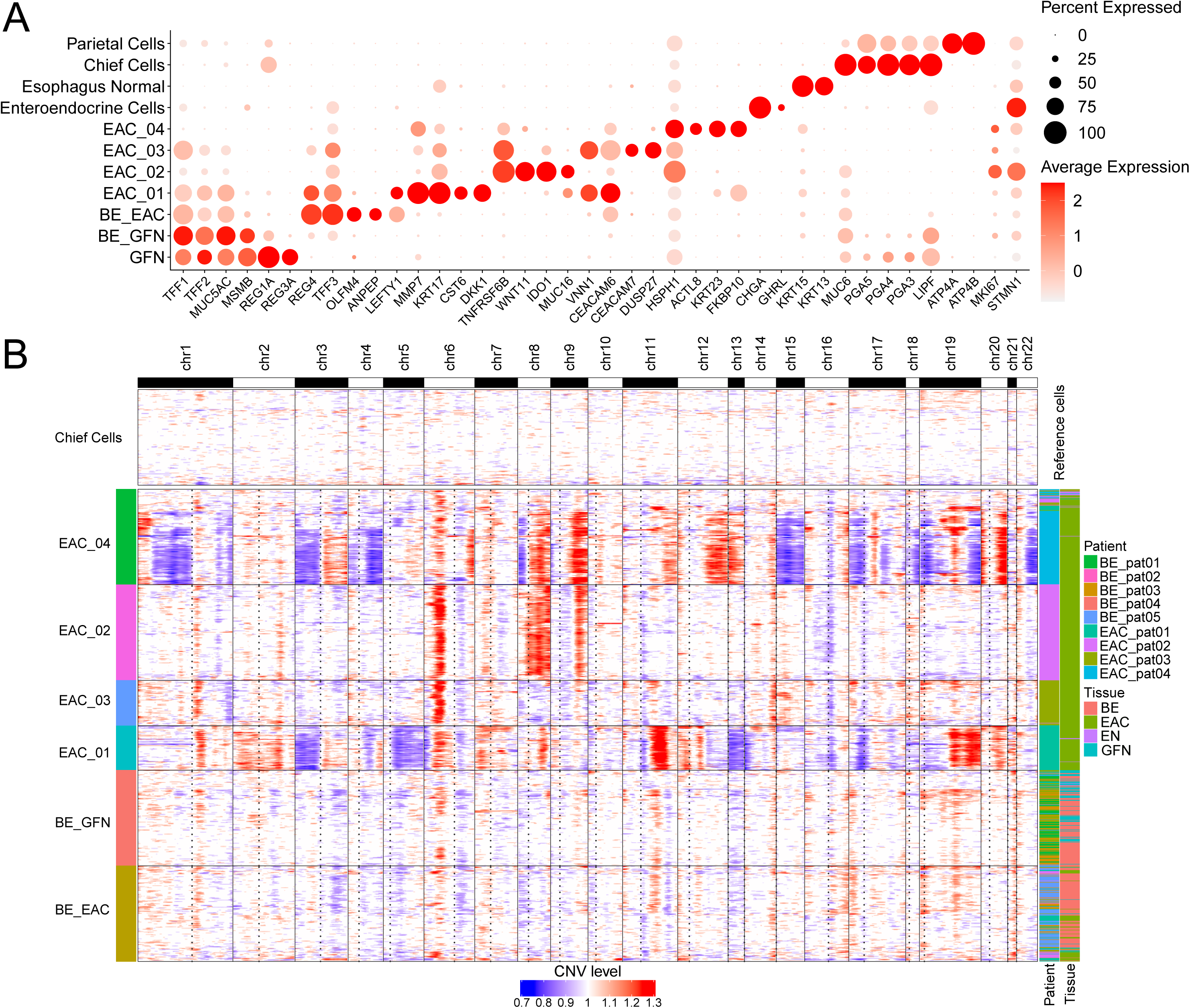
Characterization of epithelial cell types and identification of EAC-typical SCNA profiles and relative similarities. **A** Top marker expression (x-axis) per epithelial cluster (y-axis). **B** On the x-axis, genes are ordered regarding their physical location in the genome. Full lines indicate the chromosomes, dashed lines the approximate position of the centromere. On the y-axis, single cells from selected epithelial clusters are shown. The SCNA level is calculated relative to chief cells which were used as reference.

To discern tumor cells from benign cells, we estimated cluster-specific somatic copy number alterations (SCNAs) based on single cell transcriptomic data as described by Tirosh et al. ^10^ using the R package *inferCNV*. We found that tumor cells bore distinct patient-specific SCNA profiles in contrast to benign cells (**Figure 3B**). We observed for EAC-01, EAC-02, EAC-03, and EAC-04 typical EAC SCNAs ^11,12^ (**Supplementary Table 5**). Notably, EAC-02 and EAC-04 shared three large scale copy gains on 6p, 8q, and 9q. In contrast, BE-GFN and BE-EAC did not show characteristic cancer alterations.

### Functional analysis of epithelial cells reveals differences between cells from BE and across EAC tumors

To get a better understanding of the functional properties of the epithelial cell types and subgroups we used selected cell type-specific gene-sets from Busslinger et al. ^13^ (**Figure 4A**, upper heatmap panel) as well as other gene sets regarding benign and malignant esophageal cells or samples (**Figure 4A**, lower heatmap panel) for gene-set enrichment analysis (GSEA). Unsupervised clustering of the results showed a qualitative contrast between cells from EAC-01/EAC-03 and EAC-02/EAC-04. EAC-01/EAC-03 demonstrated an enrichment of terms related to gastric pit cells, esophageal late suprabasal cells, and esophageal *KRT6B+* secretory progenitor cells from Gao et al. ^14^ (**Figure 4A**). In contrast, EAC-02/EAC-04 that were characterized as highly proliferative cells (**Figure 2D**) exhibited an exclusive enrichment of esophageal early suprabasal cells (**Figure 4A**).

**Figure 4.**
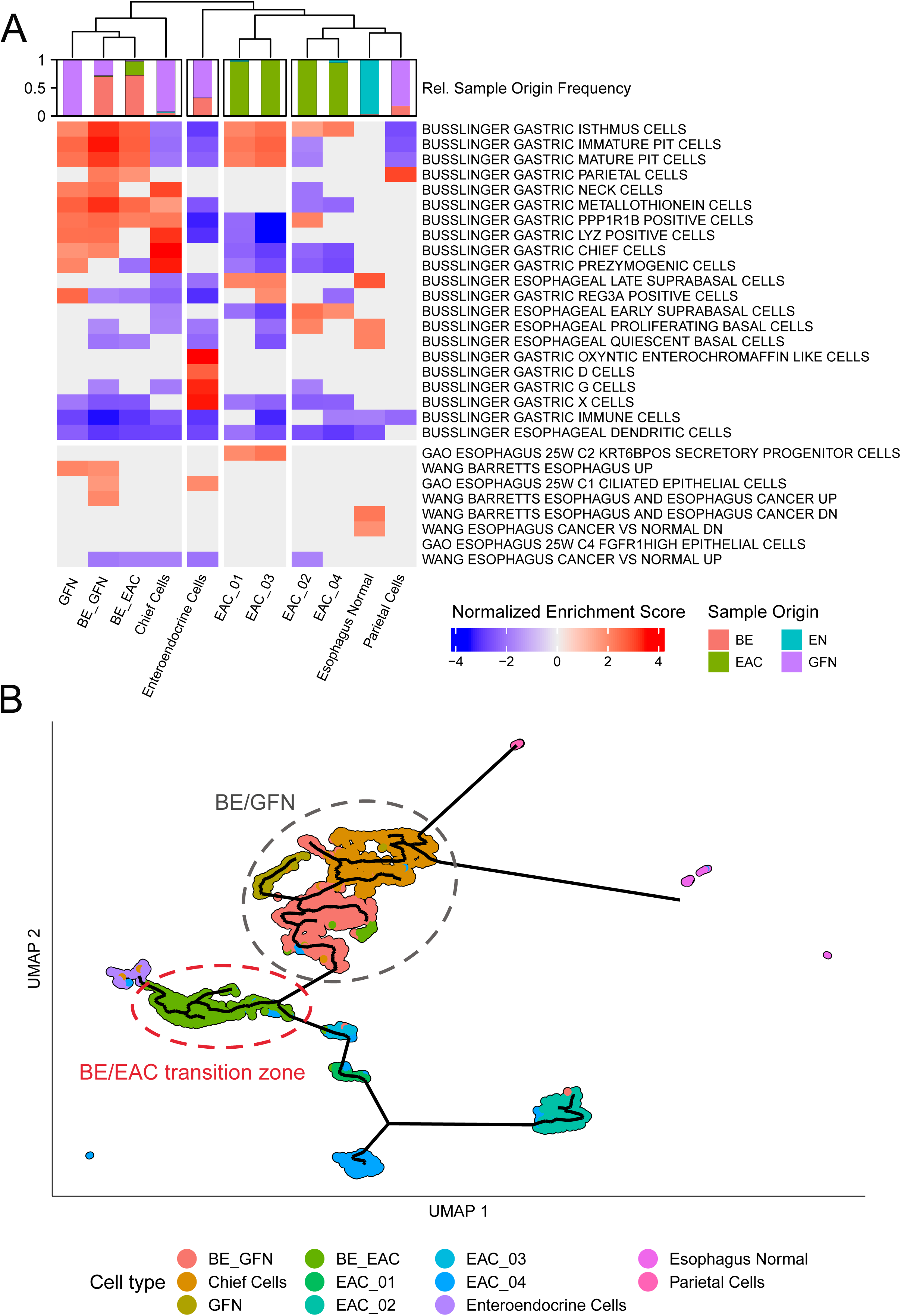
Characterization of epithelial cells shows gastric features for BE-EAC. GSEA of epithelial clusters. Enrichment of gene sets from Busslinger et al. ^13^ with relationship to BE or the esophagus were tested for enrichment in cell types (columns). Top row represents the origin of the clinical sample for each cell type. **B** Trajectory analysis of epithelial cells. UMAP representation of epithelial cells as in Fig. 2A superimposed with the pseudotime trajectory (black lines) as a proxy for relatedness.

The three clusters BE-GFN, GFN, and BE-EAC showed a similar enrichment for the gene-sets of gastric pit cells and other cell type signatures from the stomach (**Figure 4A**) emphasizing the functional similarity of cells from BE and GFN. Notably, only BE-GFN was enriched for the gene-set *WANG BARRETTS ESOPHAGUS AND CANCER UP* from Wang et al. ^15^ (**Figure 4A**). In the latter study, these genes were upregulated in BE and EAC compared to normal esophageal tissue giving evidence for an expression-based similarity between this cell type and disease progression. In contrast, BE-EAC cells do not share these BE-characteristic features and show less features of chief and prezymogenic cells (**Supplementary Figure 6** and **Supplementary Table 6**).

Using a different approach, we analyzed the results of a GSEA regarding gene ontology gene-sets (GO). For this, the other cell types of the initial analysis were added in order to characterize epithelial cells relative to other cell types (**Supplementary Figure 7**). Regarding epithelial cancer cells, again a segregation of the tumor cells into two groups was observed. EAC-01/EAC-03 were enriched in GO terms related to cell-matrix and cell-cell adhesion, cell migration, and locomotion, while EAC-02/EAC-04 were enriched in terms related to translational processes. In accordance with the findings above (**Figure 2D**) EAC-02 also showed enrichment in G2- to M-phase transition.

### Trajectory analysis implies a route to BE and EAC

We investigated the relatedness of cell lineages by a trajectory analysis using *monocle3*. Here, the trajectory closely connected BE, GFN, and chief cells (**Figure 4B**). This is in agreement with the columnal non-squamous phenotype of BE cells and supports the model of gastric cells as the origin of BE ^8^. The trajectory connects the benign BE/GFN cell group with cells derived from GFN, BE and EAC biopsies (BE/EAC transition zone) and, finally, to the malignant cells resembling the dysplastic developmental route of EAC (**Figure 4B**). Within the BE/GFN cell group, gastric chief cells are at the beginning of the trajectory. On the other side, BE-GFN is the last cell type of the BE/GFN cell group and thus characterizes the entry into the BE/EAC transition zone (**Figure 4B**).

To understand the differences between the BE/GFN group and the BE/EAC transition zone we performed a differential expression analysis (DEA) followed by GSEA. Among the highest-ranking genes with significantly increased expression in cells of the BE/GFN group were the pepsinogen coding genes *PGA3*, *PGA4*, and *PGA5* which are mostly secreted by gastric chief cells which are part of the BE/GFN group (**Supplementary Figure 6A and Supplementary Table 6**). Furthermore, *PGC* and *LIPF* were among these top differentially expressed genes, being markers for cells commonly found in gastric pits. On the side of the BE/EAC transition zone highest ranking markers were *REG4*, *TFF3*, *PHGR1*, and claudins representing gastrointestinal and goblet cell-like features. When more specifically comparing BE-GFN cells with BE-EAC cells (instead of the broader groups) we observed a downregulation of *CLDN18* and upregulation of *CLDN3, CLDN4,* and *CLDN7* in BE-EAC cells (**Supplementary Figure 6B, C and Supplementary Table 6**) in agreement with a shift towards a more metaplastic phenotype ^16,17^.

GSEA showed translational processes to be enriched in the BE/GFN cell group (containing BE-GFN cells) putting these cells close to features of EAC-02/EAC-04 (**Supplementary Figure 6D**). Furthermore, terms related to gastric cells were among the upregulated processes in BE/GFN while the BE/EAC transition zone showed enrichments in terms related to intestinal cells suggesting a functional adaption to a more intestinal phenotype. Moreover, the BE/EAC transition zone showed enrichments in terms related to oxidative phosphorylation and cell-cell interactions supporting a metabolic shift and epithelial change. The mainly BE-derived cells (BE-EAC) of the BE/EAC transition zone lacked SCNAs (see above) and had a higher metabolic turnover (oxidative phosphorylation), thereby connecting a hallmark of cancer with a non-cancer cell type.

### Enrichment of genetic risk loci in cell type-specific gene expression profiles

We wanted to understand through which cell types and biological processes inherited genetic risk profiles lead to the development of BE and EAC. Therefore, we performed a partitioned heritability analysis ^18,19^ and constructed sets of genes that were specifically expressed in each cell type or cluster. Next, we used the method LD score regression (LDSR) on our BE and EAC GWAS data^9^ to test whether risk loci that are located in 100kb regions surrounding the most specifically expressed genes of each gene-set show an enrichment of BE and/or EAC associations. In other words, we tested whether the genes that are typically expressed in a given cell type show an enrichment for GWAS risk loci of BE and/or EAC. Of note, for this analysis it is irrelevant whether cells are derived of a patient with or without the respective risk variants since generic cell type transcriptomic profiles are analyzed and no tests for allelic associations are performed.

Overall, EAC and BE GWAS risk variants showed significant enrichments among the most specifically expressed genes (gene-sets) from epithelial cells and some specific fibroblasts, endothelial cells, and immune cells (**Figure 5A-C** and **Supplementary Table 7**). This was in contrast to GWAS risk loci for major depression disorder (MDD) ^20^, which served as negative control. The failure to find any enrichment of MDD associations validated the concept of this study.

**Figure 5.**
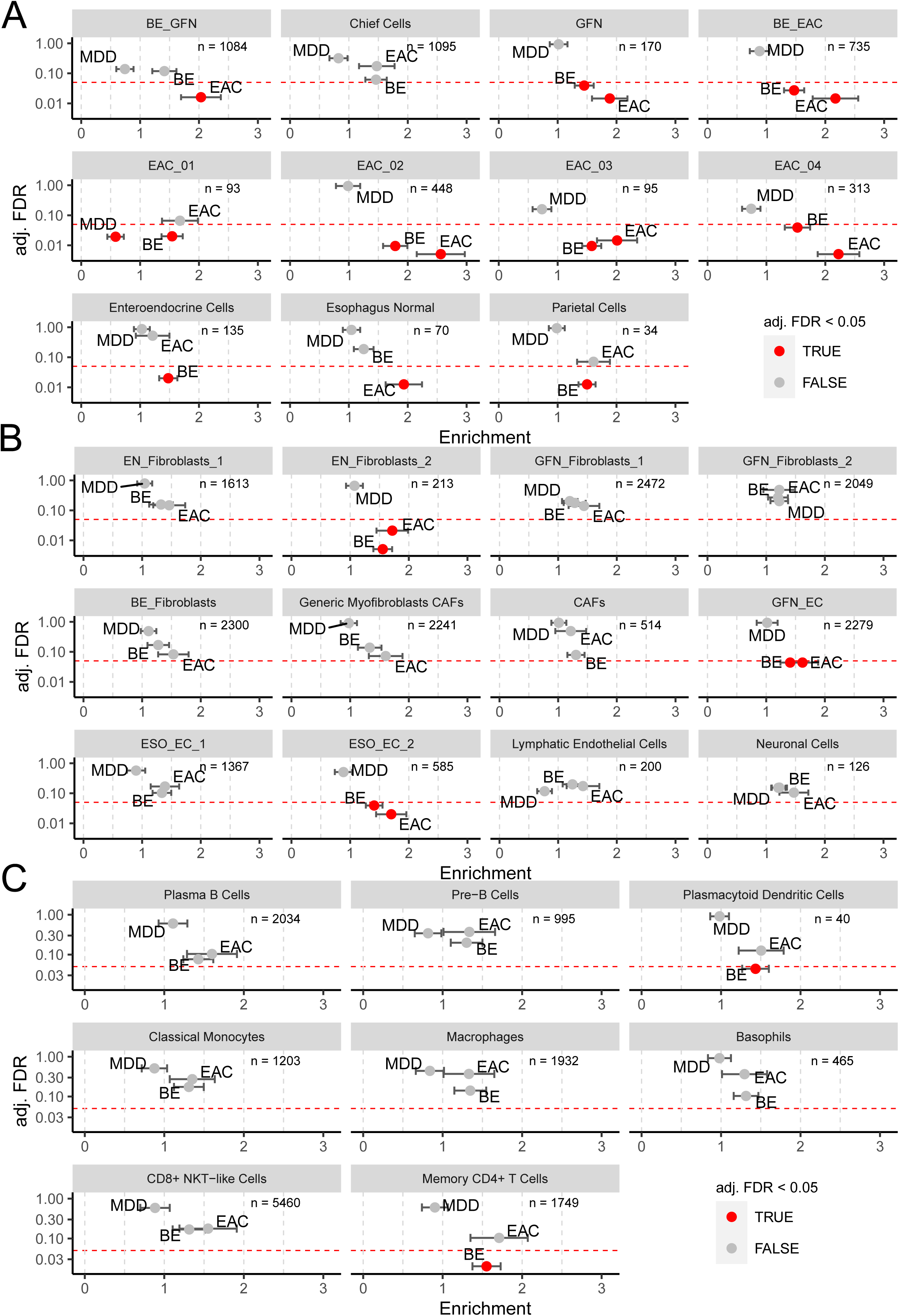
Partitioned heritability analysis of GWAS risk loci for BE and EAC applied on cell type-specific expression profiles. Fold enrichment of BE, EAC, and major depression disorder (MDD, negative control) GWAS risk loci in cell type-specific gene sets is shown on the x-axis; adjusted (adj) false discovery rate (FDR) is shown on the y-axis. **A** Epithelial cells. **B** Fibroblasts, endothelial cells (EC), esophagus normal cells (EN) and neuronal cells. **C** Immune cells. Enrichment describes the level of enrichment of GWAS risk variants within cell type-specific gene sets. Significance level with a false discovery rate (FDR) of 0.05 is indicated by a red dashed line. Significant enrichments are indicated by red data points.

In terms of significant enrichment, we observed a stronger fold-enrichment of EAC (n=10) over BE (n=5) GWAS risk loci among 31 cell types (**Figure 5A-C**). This implies that EAC development is driven to a greater extent by local cellular processes than the development of BE.

The cells of BE-EAC showed the strongest enrichment for EAC GWAS risk loci among nonmalignant cells and the enrichment was substantially weaker for BE GWAS risk loci (**Figure 5A**). This suggests that biological processes within BE-EAC cells play an important role for the malignant transformation into EAC through germline genetic risk. This adds to the evidence of a functional relationship between BE and EAC in BE-EAC cells as these non-cancer cells originate mainly from BE and to a smaller extent EAC tissue samples (**Figure 2C**). Moreover, in our trajectory analysis was BE-EAC the last cell type of the BE/EAC group that characterizes the transition to cancer cells (**Figure 4B**).

EN-fibroblasts-2, a cell type exclusively derived from normal esophagus, was enriched for EAC and BE GWAS risk loci providing evidence that fibroblasts also contribute to the etiology of EAC and BE by germline genetic risk (**Figure 5B**). In addition, endothelial cells of the ESO-EC-2 and GFN-EC type, the majority of which derived of normal esophagus and fundus, were significantly enriched for BE and EAC GWAS risk variants (**Figure 5B**) pointing at blood vessels influencing disease formation in terms of neoangiogenesis. Of the immune cells, plasmacytoid dendritic cells and CD4+ memory T cells showed enrichment exclusively for genes at BE GWAS risk loci (**Figure 5C**), indicating that processes related to the innate as well as adaptive immune system contribute to BE development through germline genetic risk.

Among the tumor cells, EAC-02 showed the strongest enrichment while EAC-01 showed no significant enrichment of EAC GWAS risk variants (**Figure 5A**). This suggests that EACs can emerge on the basis of different components of germline genetic risk.

BE-EAC characterizes the transition into EAC (**Figure 4B**) and shows substantially stronger enrichment for EAC than for BE risk loci (**Figure 5A**). These cells are only present in BE and EAC (**Figure 2C**) and represent nonmalignant epithelium (**Figure 4B**). We, thus, used a second scRNA-seq data set of BE epithelial cells from the study of Nowicki-Osuch et al. ^8^ in order to increase the cell type resolution. Multiple different epithelial cell types could be identified in this study. Applying partitioned heritability and LDSR analysis on these data, we observed enrichment of BE and/or EAC GWAS loci among genes that characterize columnar cell types. Again, MDD GWAS risk loci that served as negative control showed no enrichment and among significant enrichments, a stronger fold-enrichment among cell types was observed in EAC (n=22) compared to BE (n=3, **Figure 6**). While the enrichment was present for BE and EAC GWAS loci in columnar-undifferentiated, columnar-dividing and columnar-intermediate cell types (**Figure 6**), the enrichment in the columnar-differentiated cell type was only present for EAC GWAS loci (**Figure 6**). These findings point to cell biological processes that control the differentiation of columnar cell types as being relevant for the transformation into EAC. Marker genes of columnar-differentiated cells showed strongest enrichment in BE-EAC cells among non-cancer epithelial cells of our study (**Supplementary Figure 8**) suggesting their presence within BE-EAC cells and confirming the importance of this cell type for genetic risk-mediated EAC development. Marker genes for BE-EAC cells overlapping with columnar-differentiated cells include *REG4, CDH17, ANPEP, MUC17,* and *FABP2*.

**Figure 6.**
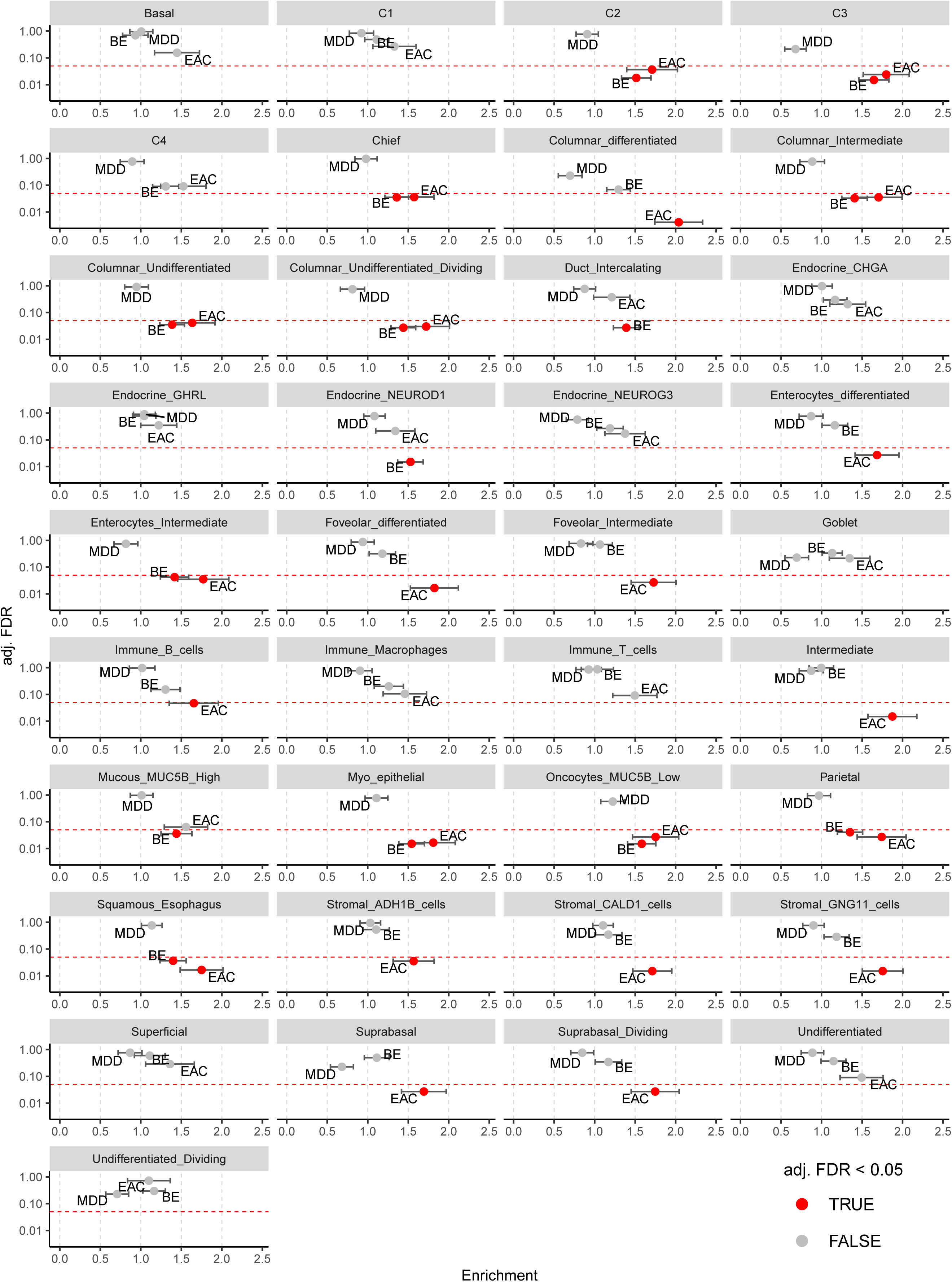
Partitioned heritability analysis of GWAS risk loci for BE and EAC applied on gastroesophageal junction cell type-specific expression profiles of Nowicki-Osuch et al. ^8^*. See* Figure 4 *for description. C1, C2, C3, and C4: KRT7^high^ (supra-)basal, submucosal gland-like cells; duct: submucosal duct;* CHGA, GHRL, NEUROD1, NEUROG3, MUC5B, ADH1B, CALD1, GNG11*: marker genes for cell types of Nowicki-Osuch et al.* ^8^.

Overall, the analysis confirms the contribution of several cell types to the development of BE and EAC and the stronger contribution of local cellular processes influenced by germline disposition for EAC compared to BE with columnar-differentiated cells within BE-EAC showing a strong contribution.

## Discussion

In contrast to earlier studies on BE or EAC scRNA-seq ^7,8,13,21,22^ our focus was to identify the cell types and biological processes that contribute to the development of BE and/or EAC through germline genetic risk (partitioned heritability ^9,18,19^).

Although the BE sample size used in the GWAS is much larger compared to EAC (11,208 BE versus 5,582 EAC patients ^9^) and thus has a higher statistical power to detect enrichment, EAC GWAS risk loci showed stronger enrichment among most of our epithelial cell types. Of all 31 cell types analyzed, 15 showed significant enrichment for EAC and/or BE risk loci. Of these, 10 showed stronger enrichment for EAC GWAS loci, whereas only 5 showed stronger enrichment for BE GWAS loci (**Figure 5**). This finding was validated in the independent scRNA-Seq of Nowicki-Osuch et al. ^8^. Twenty-two of the analyzed cell types showed stronger enrichment for EAC than BE GWAS loci and three cell types showed stronger enrichment for BE than EAC GWAS loci (**Figure 6**). Our results imply that EAC development is driven to a greater extent by local cellular processes than the development of BE. The finding confirms the results of our GWAS ^9^, where it has been shown that non-local pathomechanisms, like GERD and hiatal hernia, contribute substantially stronger to the metaplastic BE transformation than to EAC development.

Of all cell types, our data suggest that BE-EAC is of utmost importance for the malignant transformation to EAC. Our trajectory analysis showed that BE-EAC represents the last non-cancer cell type that characterizes the entry into EAC (**Figure 4B**). Accordingly, BE-EAC showed the strongest enrichment for EAC GWAS risk variants among all nonmalignant cells. This effect is stronger for EAC, as BE-EAC shows weaker enrichment for BE GWAS risk loci (**Figure 5**). BE-EAC consists of nonmalignant cells derived from BE and EAC epithelium and its benign state is in agreement with the absence of large SCNAs according to our copy number profile analysis (**Figure 3B**).

Based on the impact of BE-EAC on malignant transformation, we used the scRNA-seq data of Nowicki-Osuch et al. ^8^ and applied partitioned heritability/LDSR analyses. In this study, the BE epithelium was analyzed in more detail and led to the identification of a diversity of different epithelial cell types. We found that columnar-undifferentiated, columnar-dividing and columnar-intermediate cell types showed significant enrichment for BE and EAC GWAS risk loci. In contrast, the columnar-differentiated cell type exclusively showed enrichment for EAC GWAS risk loci (**Figure 6**). It is known that columnar epithelial cells develop from undifferentiated via dividing and intermediate to differentiated cells. Among these cell types, the undifferentiated columnar cells constitute the origin of EAC according to Nowicki-Osuch et al. ^8^. Our data confirm that undifferentiated columnar cells are of relevance in EAC development, but show that germline genetic risk factors that control all steps of the cellular differentiation of the columnar BE epithelium contribute to EAC. This also includes the last step of columnar differentiation into differentiated cells that showed the strongest enrichment of EAC GWAS risk loci in the BE epithelium scRNA-Seq data set.

Genes that are characteristic for columnar differentiated and BE-EAC cells include *REG4, CDH17, ANPEP, MUC17,* and *FABP2*. *Regenerating islet-derived family member 4* (*REG4*) is a member of the calcium-dependent lectin gene family. It is a marker for deep crypt secretory cells of the small and large intestine that provide secretory support for stem cells at the bottom of crypts ^23^. Upregulation of *REG4* has been reported for gastrointestinal cancers where its dysregulation has been connected with proliferation, invasion, and drug resistance ^24,25^. *REG4* promotes invasion, proliferation, tumor growth but also migration of gastric cancer cells ^26,27^. *Cadherin-17* (*CDH17*) is a marker of differentiation in intestinal cells and promotes cell adhesion and proliferation in colorectal cancer ^28^. *ANPEP* (*alanyl aminopeptidase formerly CD13*) is a surface marker of mature zymogenic chief cells in the gastric epithelium ^29^ and is downregulated in colorectal cancer compared to normal colon tissue ^30^. *Mucin 17* (*MUC17*) encodes for a transmembrane glycoprotein with a large extracellular domain. It is expressed in intestinal cells ^31^ and assumed to protect the intestinal epithelium from microbial invasion. *Fatty acid binding protein 2* (*FABP2*) is involved in uptake and trafficking of lipids in the intestine ^32^. It is primarily expressed in the tips of intestinal villi ^33^. These markers describe a differentiated columnar cell type that can be found in BE and EAC tissue which contributes to EAC development through germline genetic risk variants.

Apart from the relevance of BE-EAC and columnar epithelial cells to EAC development, we found that germline genetic risk factors also influence BE and EAC through other cell types (**Figure 5**). This implies that a variety of different cellular mechanisms orchestrate BE and EAC development. A defined population of fibroblasts, namely EN-fibroblasts-2, showed enrichment for BE and EAC GWAS risk variants. Whereas the role of cancer-associated fibroblasts (CAFs) on treatment response and the formation of metastases is well documented ^34^, the influence of normal fibroblasts on the development of cancer is largely unknown and provides a new area for research. In addition, two endothelial cell types, ESO-EC-2 and GFN-EC, were significantly enriched for BE and EAC GWAS risk variants pointing at blood vessels being crucial for disease formation, presumably through neoangiogenesis. The distinct enrichment of BE GWAS risk variants in plasmacytoid dendritic cells and in memory CD4+ T cells further emphasizes the complexity of BE etiology. It suggests that crosstalk between the innate immune system through interferon γ secretion by plasmacytoid dendritic cells and the adaptive immune system through storage of successful T cell receptors (memory T cells) upon antigen-triggered activation are involved processes in BE etiology. It is well known that the adaptive immune system is relevant for the elimination of (pre-) malignant cells carrying neoantigens. Our data thus suggest that germline genetic factors influence the adaptive immune system and may impact the immune escape rate of mutation carrying BE clones to eventually grow. Since it has been shown that BE already contains cancer-like mutations ^35^, it is plausible that neoepitopes at this benign stage can already be targets for immune response, involving memory T cells.

Finally, the partitioned heritability analysis showed that tumor cells are enriched for EAC GWAS risk variants to a different, tumor-dependent degree, ranging from high (EAC-02, EAC-04) to low enrichment (EAC-01). This finding is quite interesting and indicates that germline genetic factors are of different relevance for the development of individual EACs.

Overall, our study demonstrates that certain local cell types contribute to BE and EAC development through germline predisposition. These effects are substantially stronger in the development of EAC than BE, where non-local factors such as GERD and hiatal hernias are also involved. We identified BE-EAC as a cell type of pivotal relevance for EAC development with a phenotype that can be found in BE epithelium. Here, cell biological processes that control the differentiation of columnar epithelial cells seem to be mainly involved in EAC development. Thus, the diagnostic use of markers that characterize the differentiation of columnar epithelial cells should be explored in future for EAC prediction.

## Acknowledgments

The study was supported by the Wilhelm Sander-Foundation with grant 2020.119.1, the German Research Foundation (DFG) with grants 418074181, 446411360, CRC 1310 C4, and the Federal Ministry of Education and Research (BMBF) with grant CompL DeepInsight 031L0267B. Fig. 1A was created with BioRender.com. We thank the patients who participated in the study.

## Author contributions

JS and AMH designed and coordinated the study. PSP, SH, CJ, CJB, YZ and S-HC collected clinical samples. PSP, SH, RB, AQ and S-HC obtained and analyzed clinical data. PSP and SH dissociated tissue samples. MF performed NGS. MCW and A-SG performed bioinformatics analyses. MCW, A-SG, PD, DH, CP and CM performed data analysis. MCW, A-SG, RT, YZ, RCF, RB, AQ, IG, CM, JS and AMH interpreted the data. MCW, JS and AMH drafted the manuscript.

## Declarations

### Ethics approval and consent to participate

The study was approved by the institutional review board of the University of Cologne (18-274), follows the principles of the convention of Helsinki, and informed consent was obtained from all participating patients.

### Availability of data and materials

The scRNA-seq data has been deposited at the European Genome-Phenome Archive (EGA) with accession number EGAS50000000530. GWAS data was previously published ^9^ with the dataset for BEACON being available in dbGAP and the dataset of the Cambridge cohort being available via EDAM. The scRNA-seq data of Nowicki-Osuch and colleagues available from the European Genome-phenome Archive (EGA), www.esophaguscancercellatlas.org, and the HCA project as indicated in ^8^.

## Methods

### Patient cohort

Samples from two groups of patients were analyzed, one comprising five patients with pre-malignant metaplastic BE and one with four EAC patients. All patients were males of European descent. During diagnostic esophagogastroscopy biopsies were taken from the metaplastic or cancer tissue along with biopsies from normal esophageal (EN) epithelium (>5 cm proximal of BE/EAC) and normal gastric samples from the fundus (GFN).

### Sample processing

Fresh tissue samples from endoscopies, taken either from the site of the pathologies or at a distance of 5 cm apart for normal tissue, were dissociated as described earlier ^36^. In brief, the tissue was minced to small pieces, disrupted in a C-tube used with the GentleMACS Dissociator (Miletnyi Biotec) combined with enzymatic digestion with DNAse I (500 *U* ⋅ *mL*^−1^; AppliChem PanReac), collagenase IV (320 *U* ⋅ *mL*^−1^; Thermo Fisher Scientific), and dispase II (2 *U* ⋅ *mL*^−1^; Sigma-Aldrich), filtered through a 100 *µm* cell strainer, collected and resuspended in 60% RPMI-1640 medium (Thermo Fisher Scientific), 30% FBS (Capricorn Scientific), and 10% dimethyl sulfoxide (DMSO) (Sigma-Aldrich) for freezing at −80°C. After 24 hours samples were transferred to liquid nitrogen until fluorescence activated cell sorting (FACS). FACS was conducted prior to sequencing.

The samples from patients with BE were processed in the following way: Single-cell suspensions were stained with PE/Cy7-conjugated antibodies against CD45 (Biolegend) as a leukocyte marker and with propidium iodide (Thermo Fisher Scientific) to distinguish live and dead cells according to the manufacturer’s specifications. For the cells from EAC samples, dissociated single-cell solutions were prepared for Cellular Indexing of Transcriptomes and Epitopes by Sequencing (CITE-Seq) analysis. After thawing, cells were prepared with the kit for the cell surface protein labeling for single cell RNA sequencing (10x-Genomics) according to the manufacturer’s instructions. First, cells were labeled with 2 µl preconjugated antibodies per 100 µl total volume. Antibodies (category: TotalSeq™-B) against CD45 (order nr: 304066, barcode sequence TGCAATTACCCGGAT), CD90 (order nr: 328147, barcode GCATTGTACGATTCA), CD326 (order nr: 32429, barcode TTCCGAGCAAGTA), and CD31 (order nr: 303145, barcode ACCTTTATGCCACGG) were purchased from BioLegends and each labeled with specific barcode sequences. To avoid unspecific binding, Human TruStain FcX™ (Fc Receptor Blocking Solution) (BioLegends) (5 µl per 100 µl total volume) was added. In the next step, cells were washed to eradicate excessive antibodies. Afterwards, single-cell suspensions were stained with Hoechst (staining of all cells) and propidium iodide (staining of dead cells) (Thermo Fisher Scientific) to distinguish live and dead cells according to the manufacturer’s specifications. FACS was performed using a 100-*µm* nozzle. For each sample, 10,000 living cells were sorted. For samples with excessive leukocyte presence, CD45-positive cells were restricted to maximum 25 % of total cells. Collected single cells were placed on ice and further processed by the Cologne Center for Genomics (CCG, Cologne) for 3’ single-cell RNA-sequencing with the 10x Genomics’ scRNA-seq technology (Chromium Single Cell 3’ Solution) according to the manufacturer’s specifications. Targeted cell recovery was aiming at 3,000 cells per sample.

### Sequencing and raw data processing

The samples were processed using the 10x Genomics Chromium platform and sequenced on a NovaSeq 6000 (Illumina) with Illumina 3’ v3.1-paired end chemistry, 29-10-10-89 bp aiming at 25,000 read pairs per cell. Subsequently, the raw data was processed using the 10x Genomics software Cell Ranger v6.1.2 ^37^. Seurat V4 was then used for further handling of the data ^38^. Procedure of quality control, batch correction, and normalization are described in **Supplementary Methods.**

### Quality control of the single cell RNA-sequencing data

The output matrices of Cell Ranger were loaded into Seurat using R. Cells with at least 1,000 different sequenced genes were retained from BE patients. For cells from EAC patients and patient BE_pat02 the threshold was lowered to 500 to capture a wider spectrum of (tumor) cells. The quality of cells was inspected regarding the total number of counts per cell, the number of different genes, and the fraction of mitochondrial genes. Three median absolute deviations were allowed for the first two parameters to account for very low-quality cells and to reduce the chance to capture doublets. For the fraction of mitochondrial genes, a rather high initial threshold of 30% was chosen allowing cells with naturally higher fractions of mitochondrial mRNA to be retained. After initial clustering, the top 15% of the cells per cell type regarding that parameter were discarded to set a cell type dependent threshold.

### Batch correction, normalization, clustering, and visualization

To account for technical artifacts due to processing of cells in different batches the Canonical Correlation Analysis method included natively into the Seurat package was used before clustering and visualization ^39^. True batches, i.e. samples that were sequenced together, were merged without using a batch correction method. The data were normalized using Seurats “LogNormalize” function. Using the function *FindVariableFeatures,* 6,000 features were identified using the ‘vst’ option. Principal component (PC) analysis was performed and the number of retained PCs was determined by the point at which the smoothed difference between the accounted variability of two adjacent points was lower than 0.02 using a rolling average of two additional points in both directions. To find clusters the recommended functions *FindNeighbors* using a KNN graph based on the euclidean distance in PCA space and analyzing the Jaccard similarity and *FindClusters* which employs the Louvain algorithm were used ^38^. The *UMAP* algorithm was used to find a two-dimensional projection of the data.

### Differential expression analysis and cell type annotation

*FindAllMarkers* was used to find cluster-specific marker genes. Therefore, a minimal percentage of cells of 30% expressing a tested gene within a given cluster and a minimal logFC threshold of 0.1 were chosen as parameters. To minimize putative batch effects, we included the batch as a latent variable into the model. *MAST* was incorporated in the model by choosing the respective option ^40^. To the results, the ratio between the fraction of cells within a given cluster expressing a given gene and the fraction in all other clusters was calculated to find a measure of cluster-specificity (pct.ratio). As marker genes, the genes that had the highest logFC between clusters and the highest pct.ratio were considered. Typical markers were found for the initial cell type annotation. *PanglaoDB* was used for a semi-automatic way of cell type annotation ^41^. The immune cells were annotated following the standard workflow of *ScType* ^42^.

### Cell-cycle scoring

Using the method of Tirosh et al. ^10^, a correlation between the expression profiles of single cells and cell cycle expression signatures were calculated to assign the most probable cell cycle phase out of G1, S and G2/M. The number of cells per cluster being assigned to the S and G2/M phase were divided by the number of G1 cells in that cluster to assign the relative proliferation index.

### Gene set enrichment analysis

Gene set enrichment analysis (GSEA) was performed on gene lists obtained by the Seurat functions *FindAllMarkers* or *FindMarkers* using a logFC threshold of 0.05 and a minimum percentage of cells expressing a given gene in the tested cluster of 0.3. The used model was *MAST*. The R package *fgsea* was used to run the analysis (Korotkevich et al. bioRxiv. https://doi.org/10.1101/060012) using the databases of MSigDB (Liberzon et al., 2011, 2015; Subramanian et al., 2005).

### Inferring somatic copy number variations with *inferCNV*

The R package *inferCNV* ^10^ was used to estimate collective SCNAs using a sliding window of genes, which was set to 150. The algorithm explores expression levels of genes across the genome in comparison to a reference set of cells. The settings as recommended were used to generate the results. The order of genes follows their physical position on a given chromosome even though zero-expression genes have been discarded. The approximate physical position of the centromere for each chromosome is marked at the last included gene physically positioned before CENP-A of the respective chromosome ^43^. If that particular gene had been discarded due to zero expression, the next included gene towards the end of the chromosome was considered as a proxy.

### Trajectory analysis using *Monocle3*

For trajectory analysis we used the R package *Monocle3*, using the option close_loop = TRUE ^44^.

### Partitioned heritability

Gene-sets derived from the scRNA seq data were used to create genome annotations. Gene-sets for each of the 31 identified cell types or clusters were constructed as follows. All genes were excluded that showed no expression in any cell type or cluster. Next, a gene’s cell type or cluster expression specificity was calculated by dividing the expression of each gene in each cell type or cluster by the total expression of this gene in all cell types and clusters. Thus, the specificity ranges between 0 and 1, where 1 means that a gene is specifically expressed in a cell type or cluster, while 0 implies no expression in the respective cell type or cluster. For each cell type or cluster, the top 15% most specific genes were used to construct gene-sets. The specificity metric follows the approach of Bryois et al. ^18^. Next, we used the LDSR partitioned heritability method ^19^ and GWAS data on BE and EAC ^9^ as well as MDD ^20^. Here, we tested whether GWAS SNPs that are located in 100kb regions surrounding the most specifically expressed genes of each gene-set show an enrichment of BE and/or EAC as well as MDD associations. In order to correct for multiple testing, we applied an FDR-adjustment on the enrichment results.

### Satistics and reproducibility

Seurat V4 with the implemented statistical analysis in R was used for definition of single cell RNA-seq clusters ^38^. The differential gene expression analysis was performed using MAST ^40^ where pairs of regression coefficients through bootstrapping and an empirical Bayesian framework are used. LDSR partitioned heritability method was applied as described by Finucane and colleagues^19^.

