## Supplementary Material for "Single cell analysis of Barrett’s esophagus and carcinoma reveals cell types conferring risk via genetic predisposition"

### Supplementary Notes

#### Note 1: Mesenchymal and immune cells

To identify mesenchymal cells, we used the markers *VWF*, *PECAM1*, *PLVAP* and *CD34* for endothelial cells, *DCN*, *DPT*, *LUM* and *PDGFRA* for fibroblasts and *RGS5*, *TAGLN*, *ACTA2* and *MYL9* for myofibroblasts. Cluster 17 had a specific expression pattern and its top markers could be linked to lymphatic endothelial cells using *PanglaoDB* [1]. Immune cells were identified by the expression of *CD2* and the *CD3* subunits *CD3D* and *CD3E* for T cells, *CD79A* for all B cells and *FPR1*, *S100A8* and *S100A9* for monocytes. Cluster 20 had a specific expression of *NRXN1* and *PLP1*, identifying the origin of these cells as neuronal (for cluster nomenclature see **Supplementary Figure 2A, C**).

Fibroblasts, endothelial cells and immune cells were subjected to subanalysis, respectively. This revealed different fibroblastic states (**Supplementary Figure 4A**). A myofibroblast-like population could be discerned by the expression of *ACTA2* and a classical fibroblast-like population was marked by the expression of *POSTN*. Among the myofibroblast-like population, a cluster annotated as Generic Myofibroblasts/CAFs (cancer-associated fibroblasts) that consisted of cells from all sampled tissue origins, suggesting a cell type state that remains stable throughout the course from normal tissue via BE to EAC (**Supplementary Figure 4B**). A group of cells with high *ACTA2* expression almost exclusively contained cells from the cancer samples, which was therefore annotated as CAFs and a cluster annotated as EN_Fibroblasts_2 with cells from normal esophageal tissue. The clusters that expressed *POSTN* were homogeneous regarding sample origin (GFN_Fibroblasts_1, GFN_Fibroblasts_2, BE_Fibroblasts, EN_Fibroblasts_1 and EN_Fibroblasts_2) but consisted of different batches, suggesting that the distinct clusters did not form based on technical artifacts but rather represent genuine cell type states with differences due to the tissue origin (**Supplementary Figure 4C**). We compared the fibroblast cell populations with high *ACTA2* expression and high *POSTN* expression (**Supplementary Figure 4D**). Typical marker genes discerned the two groups, with *RGS5*, *TAGLN*, *THY1* and *FABP4* as additional typical marker for a myofibroblast-like phenotype and *DPT*, *LUM*, *DCN* and *PDGFRA* as markers for the classical fibroblast-like group. GSEA of the genes differentially expressed in the $ACTA^{high}$-group illustrated the typical functions of myofibroblasts such as *Muscle Contraction*, *Oxidative Phosphorylation* and *Cytoskeleton Organization* but not *Cell Migration* and manipulation of the ECM while the $POSTN^{high}$-group is characterized by modulation of metabolic and immune process (**Supplementary Figure 4E**).

Among the most differentially expressed genes between the generic myofibroblast-like cluster and the CAFs were genes such as *LUM*, *VCAN*, *MMP2*, *FN1* and *TIMP1* that are widely accepted as markers for CAFs. Running GSEA on the differentially expressed genes discerning these two groups, CAFs showed enrichments in *Epithelial-to-Mesenchymal Transition*, the *Matrisome* and a *Multicancer Invasiveness Signature* (data not shown).

Having neither elevated expression levels of *ACTA2* nor *POSTN*, EN_Fibroblasts_1 had special characteristics, as illustrated by GSEA of the differences between that group and the $POSTN^{high}$-group. Functions included the *Homeostasis of Metal Ions*, *Proteolysis*, *Regulation of Immune Response* and *Secretion*, while the other group had a myofibroblast-like phenotype (**Supplementary Figure 4F**).

Clustering of endothelial cells suggests tissue of origin specific characteristics with gastric fundus-derived endothelial cells separating from the rest, while clustering of immune cells was dominated by immune cell types. The immune cells were annotated using *ScType,* a tool for automated annotation of cell types, which works well for the already thoroughly characterized compartment of immune cells (**Supplementary Figure 4G, H**).

In summary, the phenotype of fibroblasts is strongly influenced by the tissue of origin even across the related tissue environments of esophageal, gastric, Barrett's mucosa, and EAC. Myofibroblasts were characterized by typical muscle and cytoskeletal features, being separated from CAFs demonstrating features such as regulation of the ECM, promoting EMT and invasiveness in cancers. Classical fibroblasts showed enrichments in metabolic and immune processes separating them from the aforementioned clusters.

#### Note 2: Identification of epithelial cell types

We annotated the clusters chief cells, parietal cells, esophagus normal and enteroendocrine cells based on the expression of marker genes (**Figure 3A**). Chief cells expressed *PGA3*, *PGA4* and *PGA5*, encoding pepsinogen which is mostly secreted by gastric chief cells [2], further supported by the high enrichment of the chief cell signature from Busslinger et al. [2] (**Figure 4A**). *ATP4A* and *ATP4B* encode for the ATP-dependent H+-pumps specific for parietal cells (Busslinger et al., 2021), *KRT13* and *KRT15* have been reported to be specific for cells from the epithelium of the healthy esophagus [2], and enteroendocrine cells were identified by the expression of *CHGA* and *GHRL* [2,3].

Among the cancer cell clusters, *MMP7*, strongly expressed in EAC_01 (**Figure 3A**), is a matrix metalloproteinase that has been reported to have various oncogenic functions, such as inhibition of cancer cell apoptosis, reduction of cell adhesion and induction of angiogeneses (as reviewed by [4]). Garalla et al. [5] found *MMP7* to have peak expression in the invasive front of EAC compared to healthy and dysplastic tissue of the esophagus. *WNT11*, a characteristic gene for EAC_02 (**Figure 3A**), promotes non-canonical Wnt signaling pathway and has been involved in various cancers, including colorectal and gastric cancer as reviewed by Katoh [6] and Chen et al. [7] influencing tumor invasion and metastasis [8].

The cluster parietal cells presented a high enrichment of the respective gene set from Busslinger et al. [2], esophagus normal cells were enriched in the gene sets regarding esophageal late suprabasal cells, and both proliferating and quiescent basal cells of the esophagus. Busslinger et al. [2] reported the characteristics of gastric *D*, *G* and *X* cells in the context of enteroendocrine cells which showed a singular enrichment in the here found group of enteroendocrine cells.

### Supplementary Tables

### Supplementary Table 1. Patient information

| Patient ID | Diagnosis | Gender | Age | Segment length (BE) [cm] | Tumor localization (EAC) | Tissues analyzed |
|---|---|---|---|---|---|---|
| BE_pat01 | BE | m | 67 | 2 | - | EN, GFN, BE |
| BE_pat02 | BE | m | 61 | 3 | - | EN, GFN, BE |
| BE_pat03 | BE | m | 46 | 2 | - | GFN, BE * |
| BE_pat04 | BE | m | 70 | 8 | - | EN, GFN, BE |
| BE_pat05 | BE | m | 63 | 3 | - | EN, GFN, BE |
| EAC_pat01 | EAC | m | 70 | - | AEG I | EN, GFN, EAC |
| EAC_pat02 | EAC | m | 76 | - | AEG I | EN, GFN, EAC |
| EAC_pat03 | EAC | m | 87 | - | AEG II | EN, GFN, EAC |
| EAC_pat04 | EAC | m | 70 | - | AEG II | EN, GFN, EAC |

* EN tissue of BE_pat03 did not result in successful preparation of NGS library

AEG - adenocarcinomas of the esophagogastric junction

AEG I - adenocarcinoma of the distal esophagus

AEG II - true adenocarcinoma of the cardia

AEG III - subcardial adenocarcinoma

**Supplementary Table 2. Distribution of cell types across samples**

Table S2 is provided as a spread sheet in a supplementary data file. It shows the number of cells assigned to the different cell types (rows) with their tissue sample origin (columns).

### Supplementary Table 3. Genes

| | |
|---|---|
| ACKR1 | Atypical chemokine receptor 1 |
| ACKR2 | Atypical chemokine receptor 2 |
| ACTA2 | alpha smooth muscle actin |
| CCl11 | chemokine (C-C motif) ligand 11 |
| CCL2 | chemokine (C-C motif) ligand 2 |
| CCL4 | chemokine (C-C motif) ligand 4 |
| CCL5 | chemokine (C-C motif) ligand 5 |
| CD79A | B-cell antigen receptor complex-associated protein alpha chain |
| CEACAM5 | Carcinoembryonic antigen-related cell adhesion molecule 5 |
| CEACAM6 | Carcinoembryonic antigen-related cell adhesion molecule 6 |
| CHGA | Chromogranin A |
| DCN | Decorin |
| DPT | Dermatopontin |
| EPCAM | Epithelial cell adhesion molecule |
| FPR1 | Formyl peptide receptor 1 |
| FRZB | Secreted frizzled-related protein 3 |
| HNF4A | Hepatocyte nuclear factor 4 alpha |
| ITLN1 | Intelectin-1 |
| KRT20 | Keratin 20 |
| KRT6B | Keratin 6B |
| KRT7 | Keratin 7 |
| LEFTY1 | Left-right determination factor 1 |
| LIPF | Gastric lipase |
| LUM | Lumican |
| MUC2 | Mucin 2 |
| MUC5AC | Mucin-5AC |
| MYL9 | Myosin regulatory light polypeptide 9 |
| NEUROG3 | Neurogenin-3 |
| NRXN1 | Neurexin-1-alpha |
| OLFM4 | Olfactomedin 4 |
| PDGFRA | Platelet-derived growth factor receptor A |
| PECAM1 | Platelet endothelial cell adhesion molecule |
| PGA3 | Pepsinogen 3 |
| PGA4 | Pepsinogen 4 |
| PGA5 | Pepsinogen 5 |
| PGC | Progastricsin |
| PHGR1 | Proline, Histidine And Glycine Rich 1 |
| PLP1 | Proteolipid protein 1 |
| PLVAP | Plasmalemma vesicle-associated protein |
| REG4 | Regenerating islet-derived protein 4 |
| RGS5 | Regulator of G-protein signaling 5 |
| S100A8 | S100 calcium-binding protein A8 |
| S100A9 | S100 calcium-binding protein A9 |
| SFRP1 | Secreted frizzled-related protein 1 |
| SFRP2 | Secreted frizzled-related protein 2 |
| SPINK4 | Serine Peptidase Inhibitor Kazal Type 4 |
| TAGLN | Transgelin |
| TFF1 | Trefoil factor 1 |
| TFF2 | Trefoil factor 2 |
| TFF3 | Trefoil factor 3 |
| VWF | Von Willebrand factor |
| WNT11 | Protein Wnt-11 |

**Supplementary Table 4. Cell type-specific expressed genes determined by differential gene expression analysis of each cell type against all other types**

Table S4 is provided as a spread sheet in a supplementary data file. Differential gene expression analysis is performed between each individual cell type against all other cells of the study. Pct.1, fraction of cells of the cell type of interest expressing the respective gene; pct.2, fraction of all other cells expressing the respective gene.

### Supplementary Table 5. Somatic copy number alterations (SCNAs) derived from scRNA-seq of tumor samples compared to recurrent SCNAs observed in EAC

| Patient/study | Copy gains | Copy losses |
|---|---|---|
| EAC-01 | 1q, 2p, 2q, 6p, 7p, 8q, 11q, 12p, 19q, 20q | 3p, 4q, 5q, 6p, 13q, 17p |
| EAC-02 | 6p, 8q, 9q | 6q, 9q |
| EAC-03 | 6p, 14q | 1q, 6q, 9q |
| EAC-04 | 1p, 3q, 6p, 6q, 8q, 9q, 11q, 12p, 12q, 17q, 20p, 20q | 1p, 1q, 3p, 4p, 4q, 8p, 15q, 17p, 18q, 19q, 22q |
| Pasello et al. [9] | 6q, 7p, 7q, 8q, 11q, 15q, 17q | 1p, 3p, 4p, 4q, 5q, 8p, 9p, 17p, 18q |
| Frankell et al. [10]* | 1p36.22, 1q21.1, 1q21.1, 1q21.3, 1q22, 3q26.2, 3q29, 6p21.1, 6p21.32, 6p21.32, 6p21.33, 6p22.1, 7q21.3, 7q22.1, 8p23.1, 8q24.13, 8q24.3, 9q34.3, 11q13.3, 11q14.1, 12p11.23, 12q15, 13q14.11, 14q11.2, 17q11.2, 17q12, 17q21.2, 18q11.2, | 1p21.2, 1p36.11, 1p36.13, 1p36.21, 21p11.1, 21q11.2, 22q11.21, 22q11.21, 22q13.32, 3q11.2, 4q35.1, 5q12.1, 5q23.1, 6p21.32, 6p21.32, 6p24.2, 7q36.2, 8p23.3, 9p13.1, 9p21.3, 9p24.1, 9q13, 10q23.31, 10q26.2, 11q25, 14q11.2, 14q32.33, 14q32.33, 15q11.1, 15q11.2, 15q13.1, 16p11.2, 16p13.3, 16q12.1, 16q23.1, 17p11.2, |

* GISTIC peaks with q value < $10^{-10}$

**Supplementary Table 6. DEG analysis BE/GFN group vs. BE/EAC transition zone related to Supplementary Figure 6**

Table S6 is provided as a spread sheet in a supplementary data file. Differential gene expression analysis is performed between cell types of Supplementary Figure 9. Pct.1, fraction of cells of the cell type of interest expressing the respective gene; pct.2, fraction of all other cells expressing the respective gene.

**Supplementary Table 7. Results of the partitioned heritability analysis**

Table S7 is provided as a spread sheet in a supplementary data file.

### Supplementary Figures

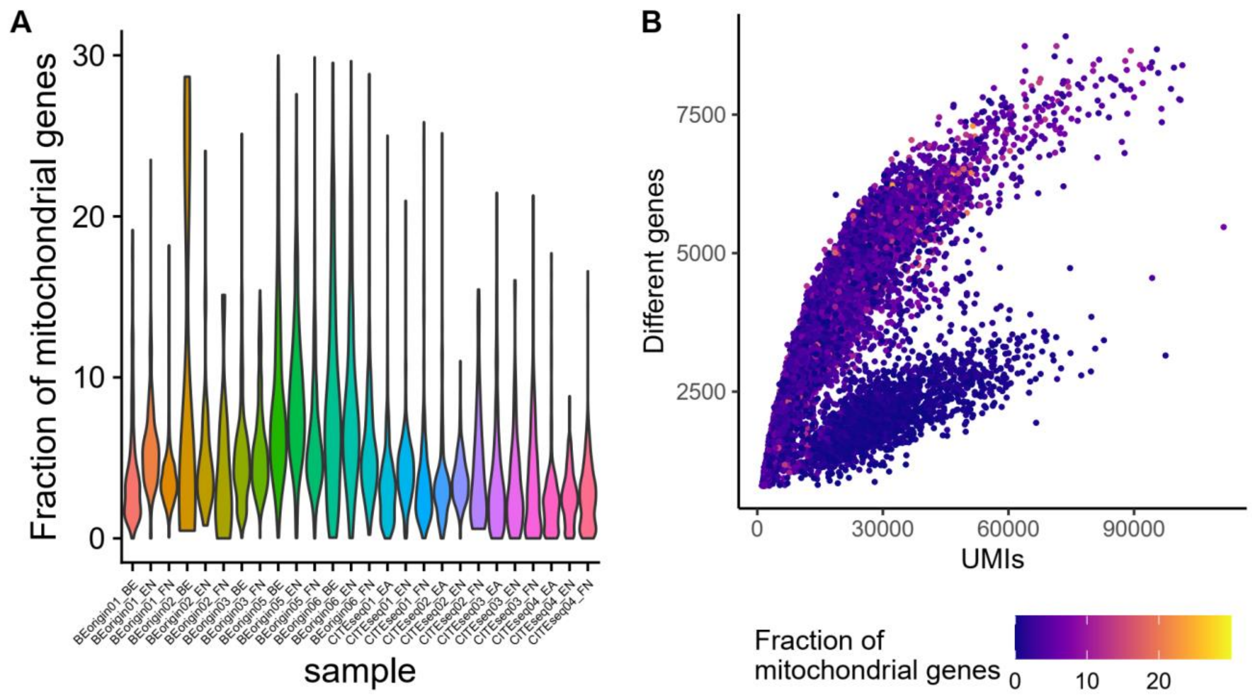

**Supplementary Figure 1. Quality Control plots. A** Percentage of mitochondrial genes regarding the total number of molecules per sample. **B** Number of different genes as a function of the total number of UMIs. Notably, the population of cells following the lower saturation curve consists of plasma B cells from different patients only.

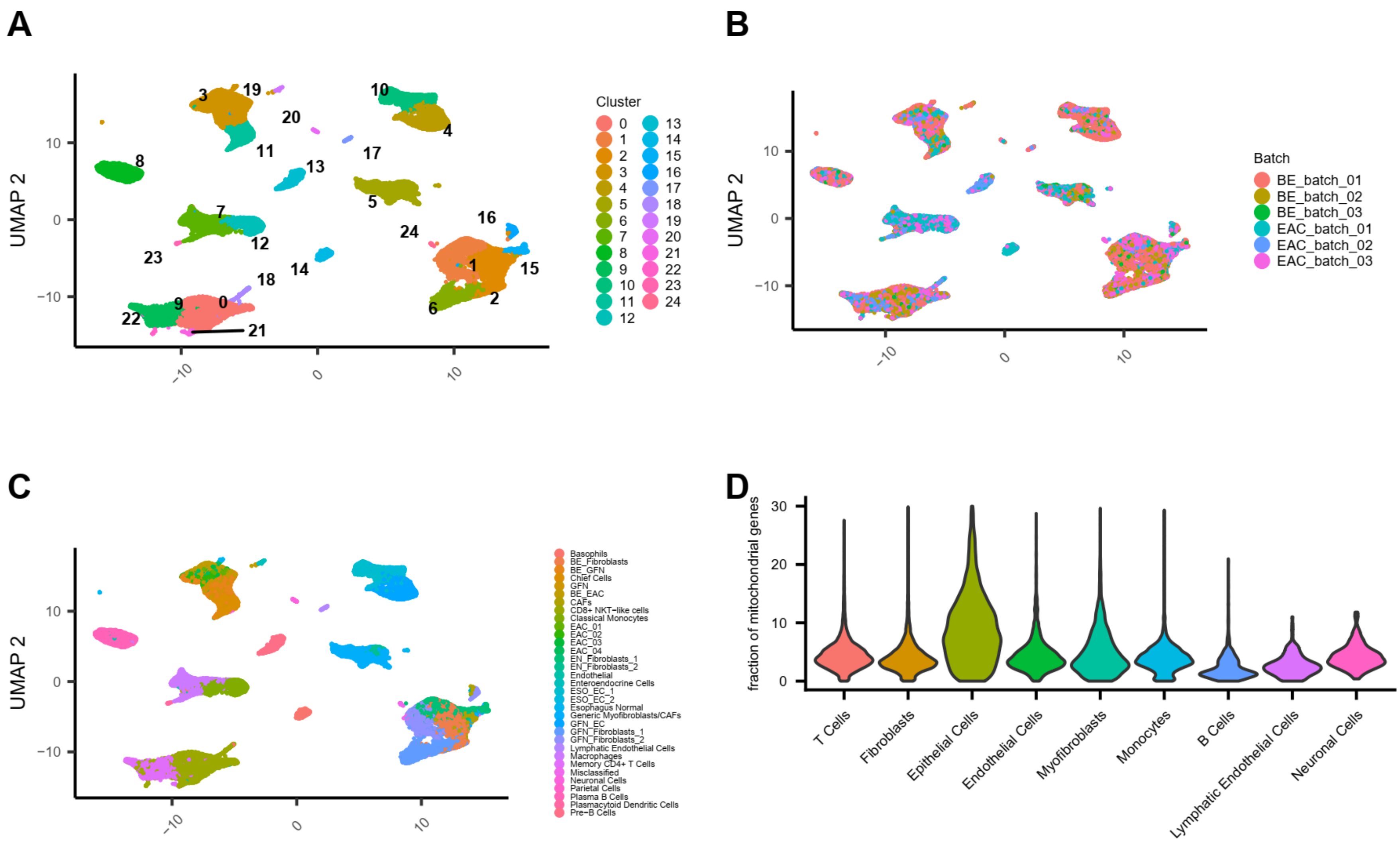

**Supplementary Figure 2.** (legend on next page)

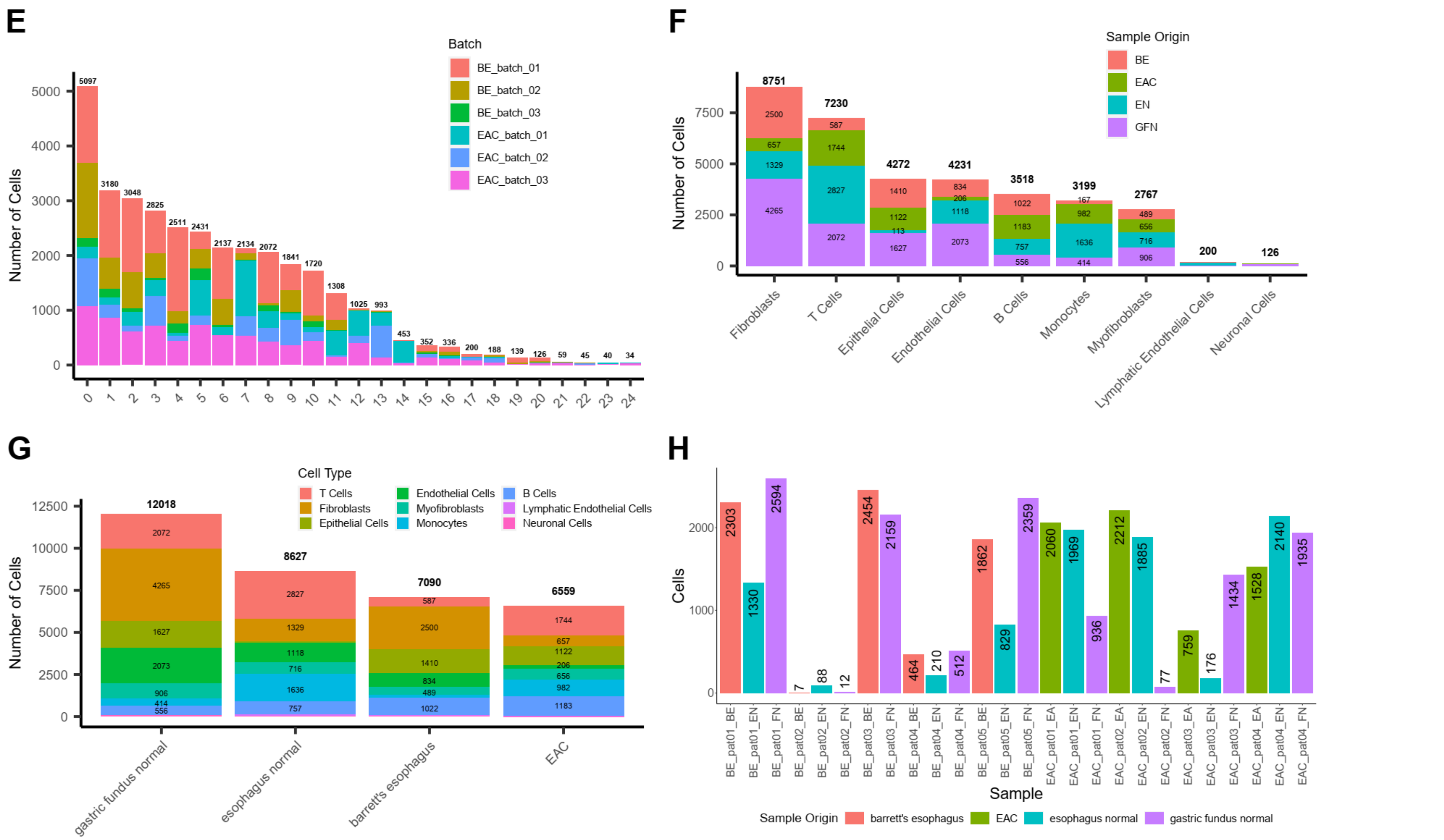

**Supplementary Figure 2. Single cell cluster characteristics. A** UMAP representation of the whole data set showing initial clusters. **B** UMAP representation with color coding for batch origin. **C** UMAP representation with color coding for subclusters. **D** Percentage of mitochondrial genes as a function of cell type family. **E** Composition of the initial clusters regarding batch origin to show batch effect control. **F** Contribution of sample origin to the cell type family. **G** Number of cells from different cell type families per sample type. **H** Cells per sample.

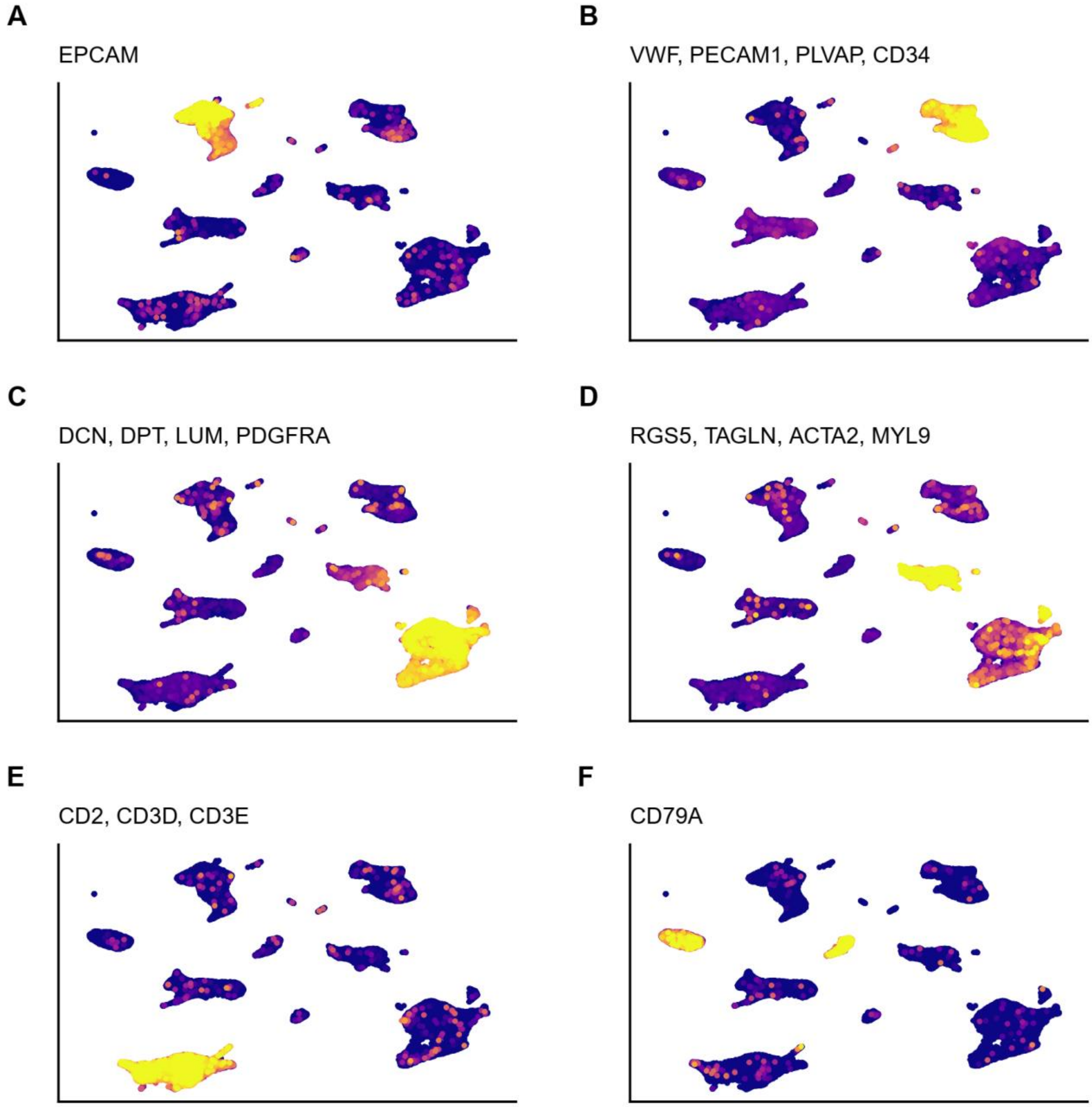

**Supplementary Figure 3. Expression of selected markers on a UMAP representation of the whole data set. A-F** Yellow color indicates high expression of the gene(s) indicated on top, *blue indicates low expression.*

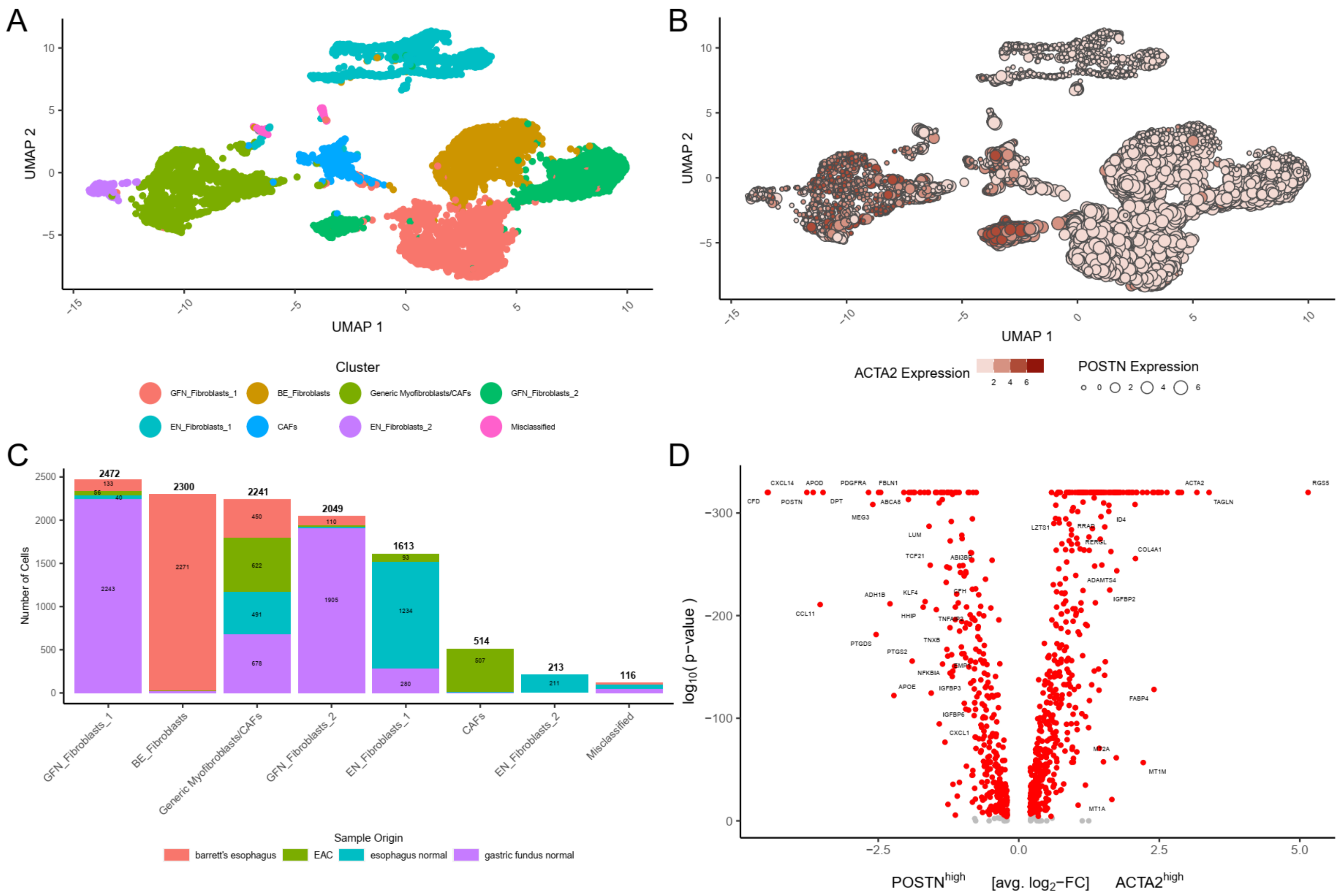

**Supplementary Figure 4** (legend on next page)

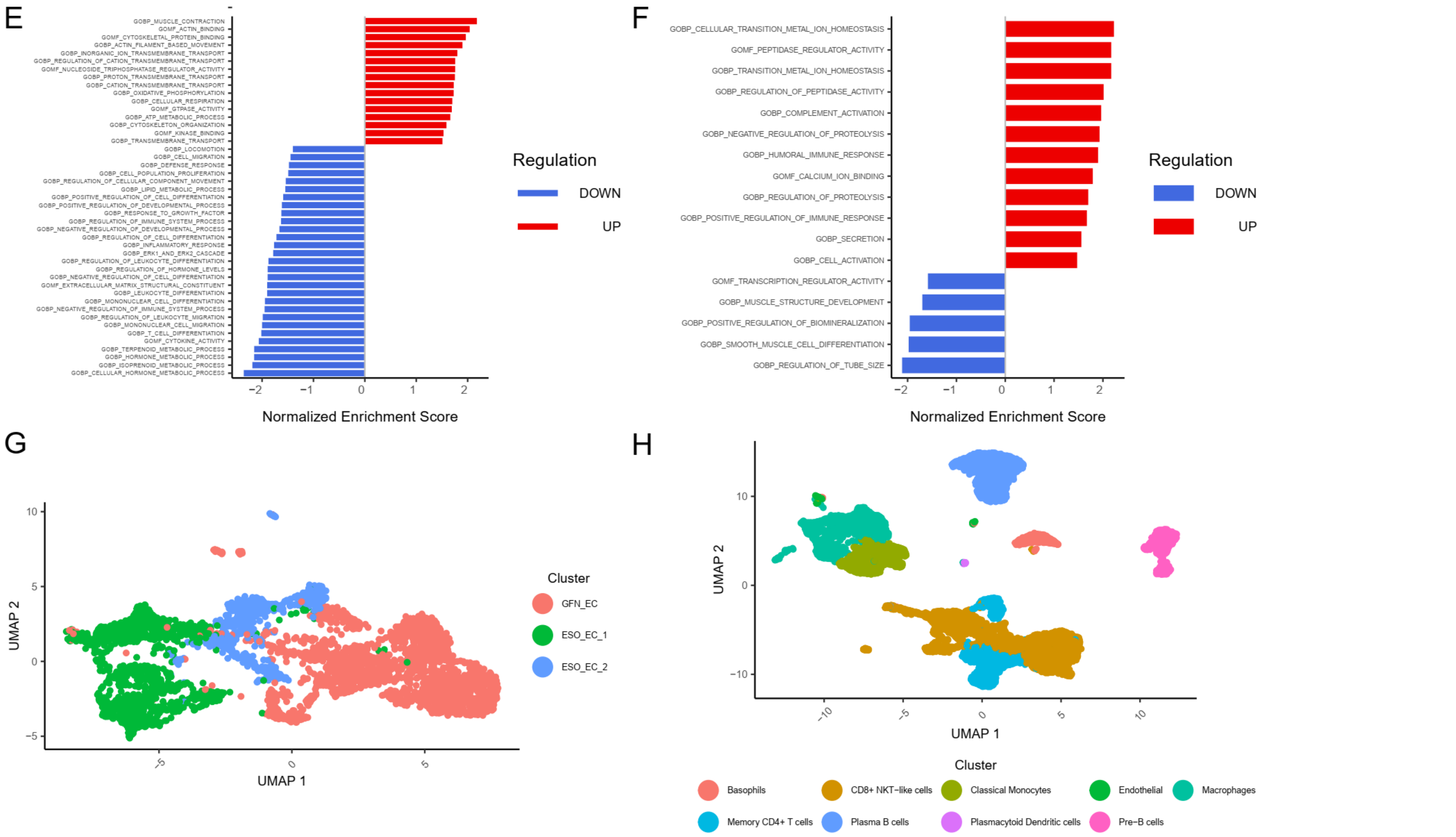

**Supplementary Figure 4. Non-epithelial cell types. A** UMAP representation of fibroblasts, myofibroblasts and CAFs after cluster annotation. **B** *ACTA2* vs. *POSTN* expression in the fibroblast-like cells. **C** Contribution of sample origins to fibroblast-like species. **D** Volcano plot of DEA of the *ACTA2*-high and the *POSTN*-high expressing clusters. **E** GSEA of myofibroblasts vs. fibroblasts. **F** GSEA of EN_Fibroblasts_1 vs. other normal fibroblasts. **G** UMAP representation of the endothelial cells after cluster annotation. **H** UMAP representation of immune cells after cluster annotation.

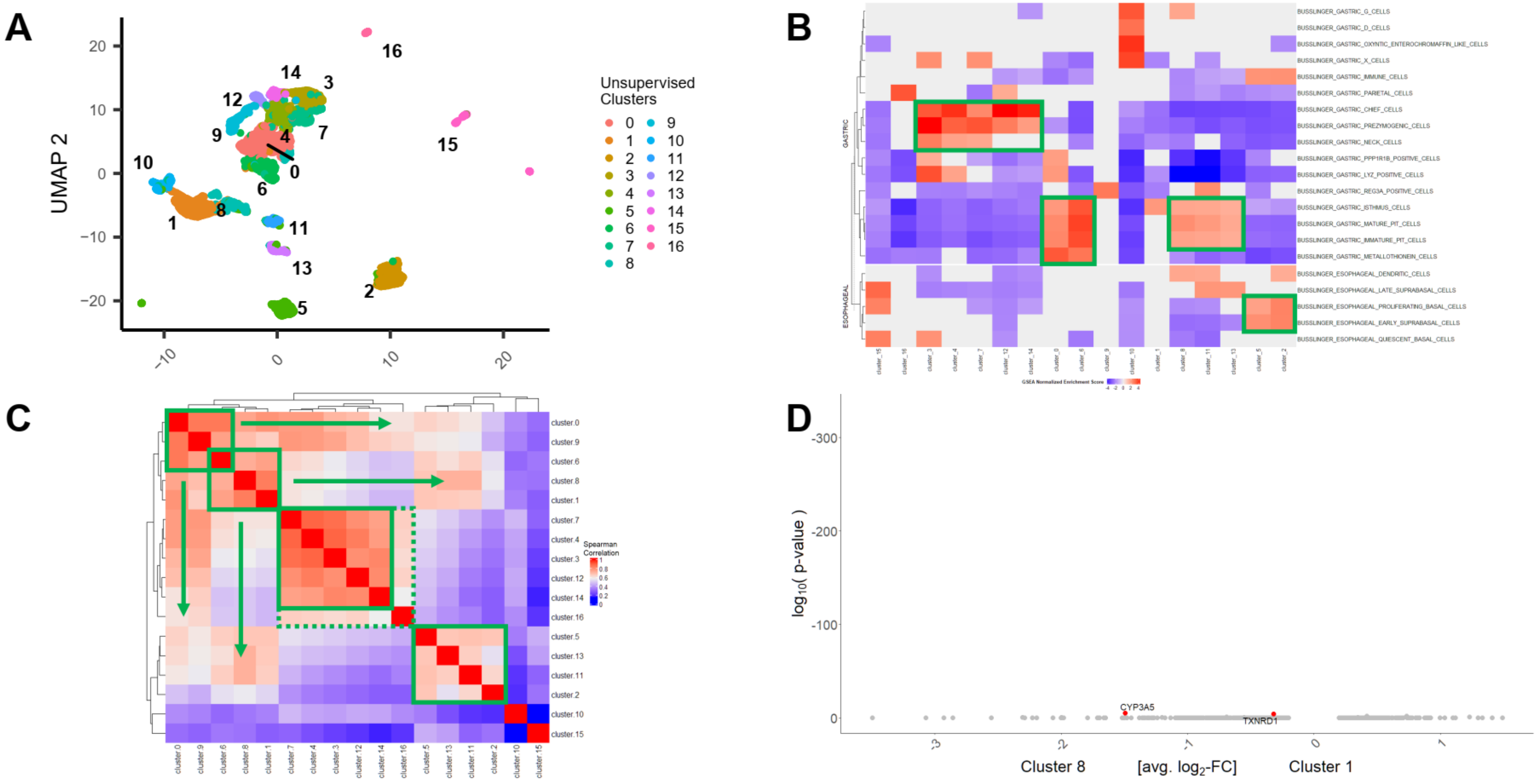

**Supplementary Figure 5** (legend on next page)

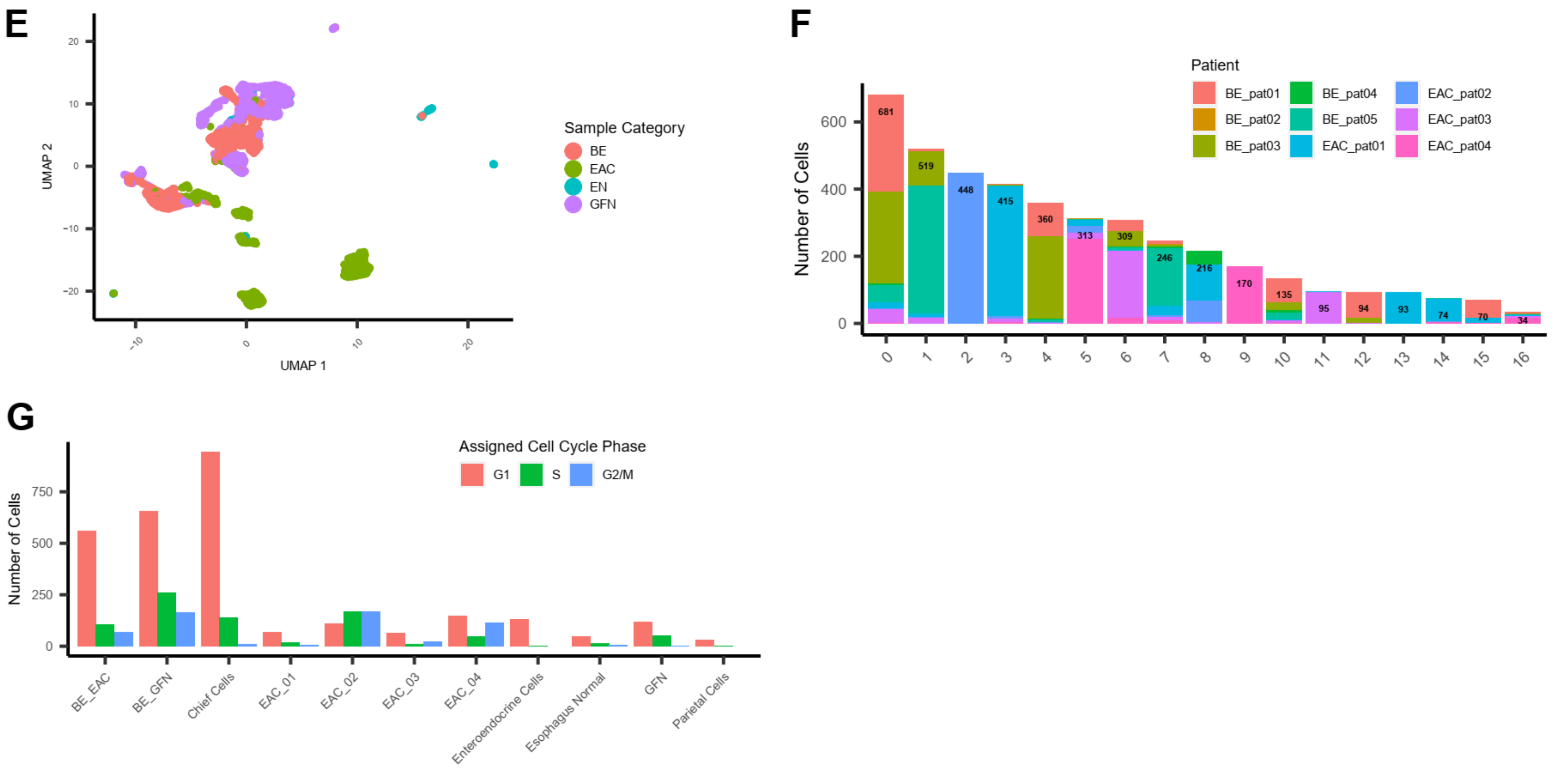

**Supplementary Figure 5. Supplementary plots of the subset of epithelial cells. A** UMAP representation of the epithelial cells with initial cluster color-coding. **B** GSEA of the initial clusters using the terms from Busslinger et al. [2]. **C** Spearman correlation matrix of initial clusters. **D** DEA of cluster 1 vs. 8 exemplifying minimal expression differences in correlated clusters. **E** UMAP representation of epithelial cells regarding sample origin. **F** Composition of initial epithelial clusters regarding patient origin. **G** Absolute number of cells in each of the cell phases from the merged epithelial clusters.

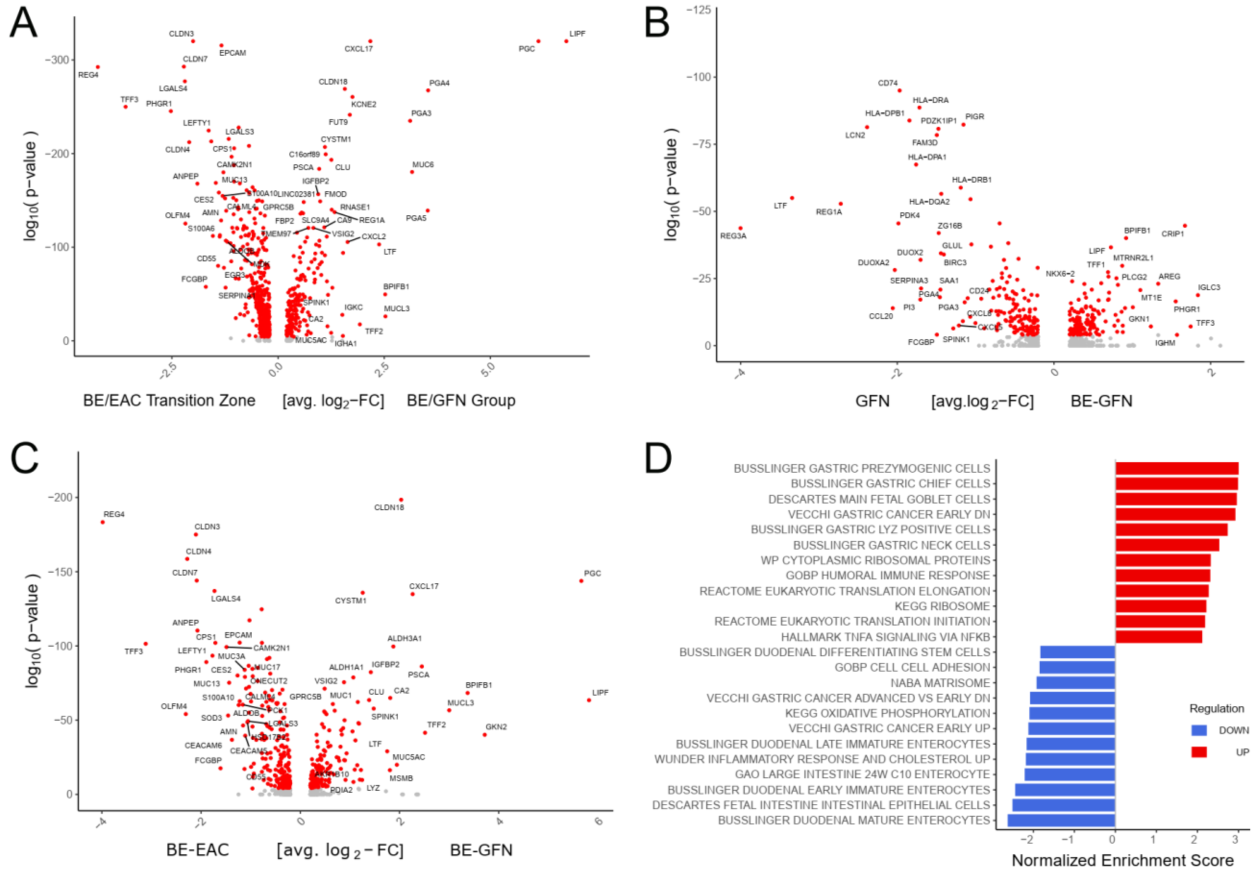

**Supplementary Figure 6. Differential expression analysis characterizing cells of the groups BE/EAC and BE/GFN. A** Volcano plot of the differential expression between cells of the BE/EAC transition zone with BE/GFN cells. **B** Volcano plot of the differential expression between GFN and BE-GFN. **C** Volcano plot of the differential expression between BE-EAC and BE-GFN. **D** GSEA of **A** with “up” referring to enrichment in BE/GFN group und “down” to BE/EAC transition zone.

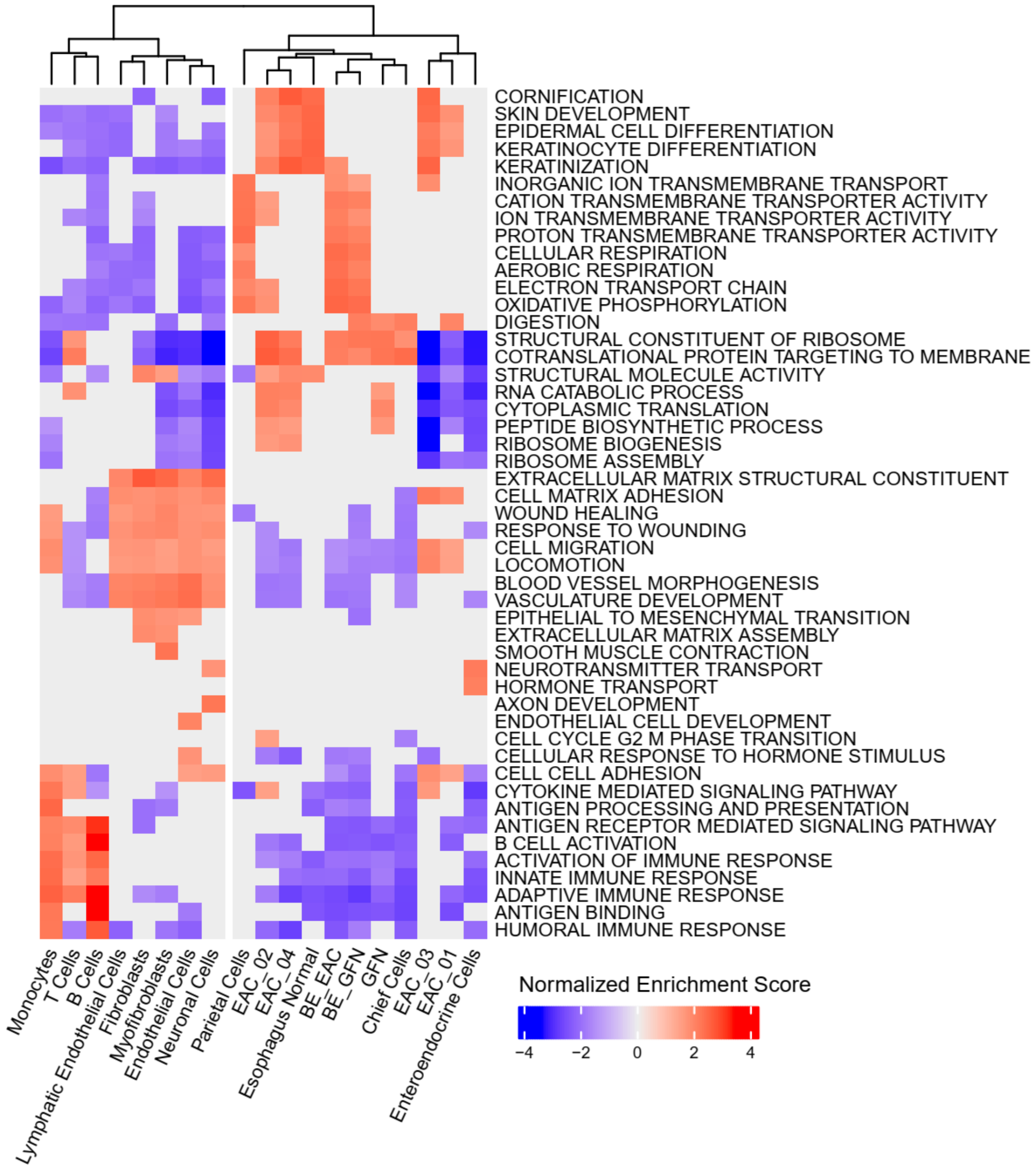

**Supplementary Figure 7. GSEA of epithelial and malignant cells in the context of non-epithelial cell types.** Selected results of GSEA of the epithelial clusters and other cell types for balance. Only GO terms are displayed.

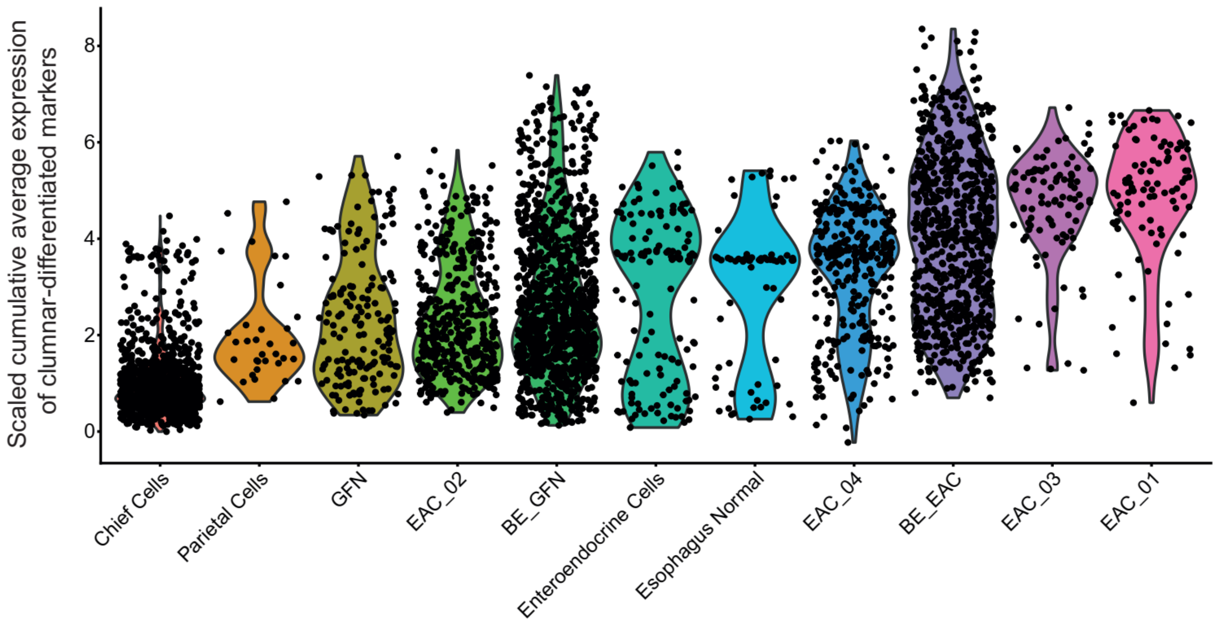

**Supplementary Figure 8. Expression of marker genes of columnar-differentiated cells of Nowicki-Osuch et al. [11].** Marker genes derived from Supp Table 7 of Nowicki-Osuch [11] were tested for expression in cells of the present study with every point representing one cell. Expression data was scaled, 0 centered with standard deviation of 1 and values represent a cumulative average per gene and cell.
